# HapSolo2: GPU-accelerated optimization for removing secondary haplotigs from diploid genome assemblies

**DOI:** 10.64898/2026.10.02.756147

**Authors:** Juan Yin, Mansi Agrawal, Prashansa Goel, Edwin A. Solares

## Abstract

**Background:** Secondary haplotigs retained in a primary assembly inflate genome size and duplicate gene content, and their removal remains a necessary curation step for heterozygous diploid genomes. HapSolo (Solares et al. 2021) treats this step as an optimization problem: it searches three pairwise alignment thresholds (percent identity, query coverage, and the ratio of aligned query to aligned reference length) to minimize a cost defined over single-copy, duplicated, and missing conserved orthologs. The original implementation depended on a separate BUSCO v3 run for every contig, on Python 2.7 and pandas in its inner loop, on a single randomized forward-walk optimizer, and on OrthoDB v9 lineage datasets that are no longer distributed.

**Results:** Here we report HapSolo2, a reimplementation that replaces the per-contig BUSCO step with miniprot-based ortholog classification against OrthoDB v9, v10, or v12.2 lineage datasets, adds steepest descent and simulated annealing to the original random forward walk, and evaluates all walkers in batched CuPy array operations on a single GPU. On an avocado assembly of 5,122 contigs scored against 1,440 orthologs, 100,000 iterations of random walk and steepest descent ran 5.6- and 7.6-fold faster on one NVIDIA RTX A6000 than on 10 CPU processes. Across 45 runs on four diploid assemblies (*Anopheles funestus*, *Persea americana*, *Vitis vinifera*, and *Amblyraja radiata*; 212 Mb to 3.2 Gb) and three OrthoDB releases, the three optimizers reached best costs within 1.8% of one another. Independent assessment with BUSCO v6.1 showed that HapSolo2 reduced duplicated orthologs from 52.0% to 5.6% in *P. americana*, from 17.9% to 10.4% in *V. vinifera*, and from 7.8% to 2.8% in *A. funestus*, with completeness decreasing by at most 1.3 percentage points. On the avocado assembly, HapSolo2 reduced the cost reached by the original HapSolo from 0.2200 to 0.1036 and raised contig N50 from 3,366 kb to 4,608 kb.

**Conclusions:** HapSolo2 makes ortholog-guided haplotig removal practical in minutes on a single workstation, independent of a BUSCO installation, and reproducible across OrthoDB releases. It is distributed as an installable Python package with a unified command-line pipeline, an assembly statistics and plotting tool, an Apptainer container, and archived copies of the legacy ortholog datasets.

## Background

Reference genomes are increasingly assembled from heterozygous individuals, due to the many species that cannot be inbred and the many phenotypes that are only visible in outbred material (Solares et al. 2021). Long-read assemblers resolve the two haplotypes of a heterozygous region into separate contigs (Chin et al. 2016; Koren et al. 2017; Cheng et al. 2021), and in regions where the haplotypes are too divergent to be recognized as alleles, both contigs are placed in the primary assembly (Roach et al. 2018; Guan et al. 2020). The result is a primary assembly that overrepresents the size and the gene content of the genome. Retained haplotigs inflate the apparent copy number of genes (Pryszcz and Gabaldón 2016; Guan et al. 2020), split read mappings between alleles so that each is covered at half the expected depth (Roach et al. 2018), and complicate downstream steps such as Hi-C scaffolding, where false joins between duplicate haplotigs produce chimeric scaffolds (Solares et al. 2021). The degree of the problem scales with heterozygosity: the avocado assembly analyzed below carries both haplotypes of most of its genome, with 52.0% of conserved orthologs duplicated.

It remains a difficult problem to identify and remove secondary haplotigs after assembly, but several solutions exist. Redundans removes the shorter of two contigs whose alignment exceeds identity and overlap thresholds (Pryszcz and Gabaldón 2016). PurgeHaplotigs and purge_dups combine read-depth histograms with self-alignment to flag contigs at haploid rather than diploid coverage (Roach et al. 2018; Guan et al. 2020). HapSolo (Solares et al. 2021) took a different approach: it performs an all-by-all alignment among contigs, defines three alignment metrics (percent identity, the proportion of the query aligned, and the ratio of aligned query to aligned reference length), and searches the space of threshold values for the triple that minimizes a cost defined by the numbers of missing, duplicated, and single-copy BUSCO genes among the retained contigs. On Chardonnay grape, the mosquito *Anopheles funestus*, and the thorny skate *Amblyraja radiata*, HapSolo reduced genome size and raised contig N50, and the reduced assemblies scaffolded into far more complete chromosomes than the unreduced assemblies did (Solares et al. 2021).

The original implementation, however, has four limitations that have become more pressing since its release. First, HapSolo required BUSCO v3.0.2 to be run separately on every contig, so that ortholog membership per contig could be tabulated before optimization. For assemblies with thousands of contigs, this preprocessing took hours to days and dominated the total run time. It also tied HapSolo to a BUSCO version that has since been superseded by versions 4 through 6, which cannot read the OrthoDB v9 datasets it depends on (Manni et al. 2021), and to those datasets themselves, which the BUSCO team no longer distributes (Zdobnov et al. 2026). Second, HapSolo was written for Python 2.7 and evaluated each candidate assembly through pandas data frame operations, which limited the number of iterations that could be afforded. Third, HapSolo offered a single optimizer, a randomized forward walk with restarts, and we noted at the time that alternative schemes such as simulated annealing were an obvious area for growth. Fourth, assembly statistics and completeness had to be computed with separate QUAST and BUSCO runs after the fact.

Here we report HapSolo2, a reimplementation of HapSolo that addresses each of these limitations while retaining the original formulation of haplotig removal as threshold optimization. HapSolo2 replaces the per-contig BUSCO runs with a single miniprot alignment of lineage ortholog proteins to the assembly followed by per-contig classification against the OrthoDB v9, v10, or v12.2 cutoffs; implements three optimizers (random forward walk, steepest descent, and simulated annealing) over a shared vectorized evaluation core; runs every walker on an NVIDIA GPU through batched CuPy array operations, with a numpy path for CPU-only systems; and packages the pipeline, an assembly statistics and plotting tool, and pinned dependencies in a pip-installable package and an Apptainer container. We evaluate HapSolo2 on four diploid assemblies that span 212 Mb to 3.2 Gb and 7.8% to 52.0% duplicated orthologs, across three OrthoDB releases and all three optimizers, assess the reduced assemblies with BUSCO v6.1 against lineages not used for training, and compare the results with those of the original HapSolo on the same assembly.

## Approach and implementation

### Overview

HapSolo2 retains the formulation of HapSolo (Solares et al. 2021). A self-alignment of the assembly yields, for every pair of overlapping contigs, three metrics: percent identity (PID), the fraction of the query contig covered by the alignment (QPct), and the ratio of aligned query length to aligned reference length (QRPct), which correspond to ID, Q and QR of the original description. For a candidate threshold triple (*p, q, r*), every contig that aligns to another contig above all three thresholds is classed as secondary and removed, and the remaining contigs form a candidate primary assembly. Each candidate is scored by the numbers of single-copy (S), duplicated (D), fragmented (F) and missing (M) conserved orthologs among the retained contigs, and an optimizer searches for the triple that minimizes a weighted cost of these counts. Only ortholog membership per contig is needed to score a candidate, so the ortholog search is run once before optimization, and each optimizer iteration reduces to set and count operations that vectorize on a CPU and on a GPU (Figure 1).

**Figure 1:**
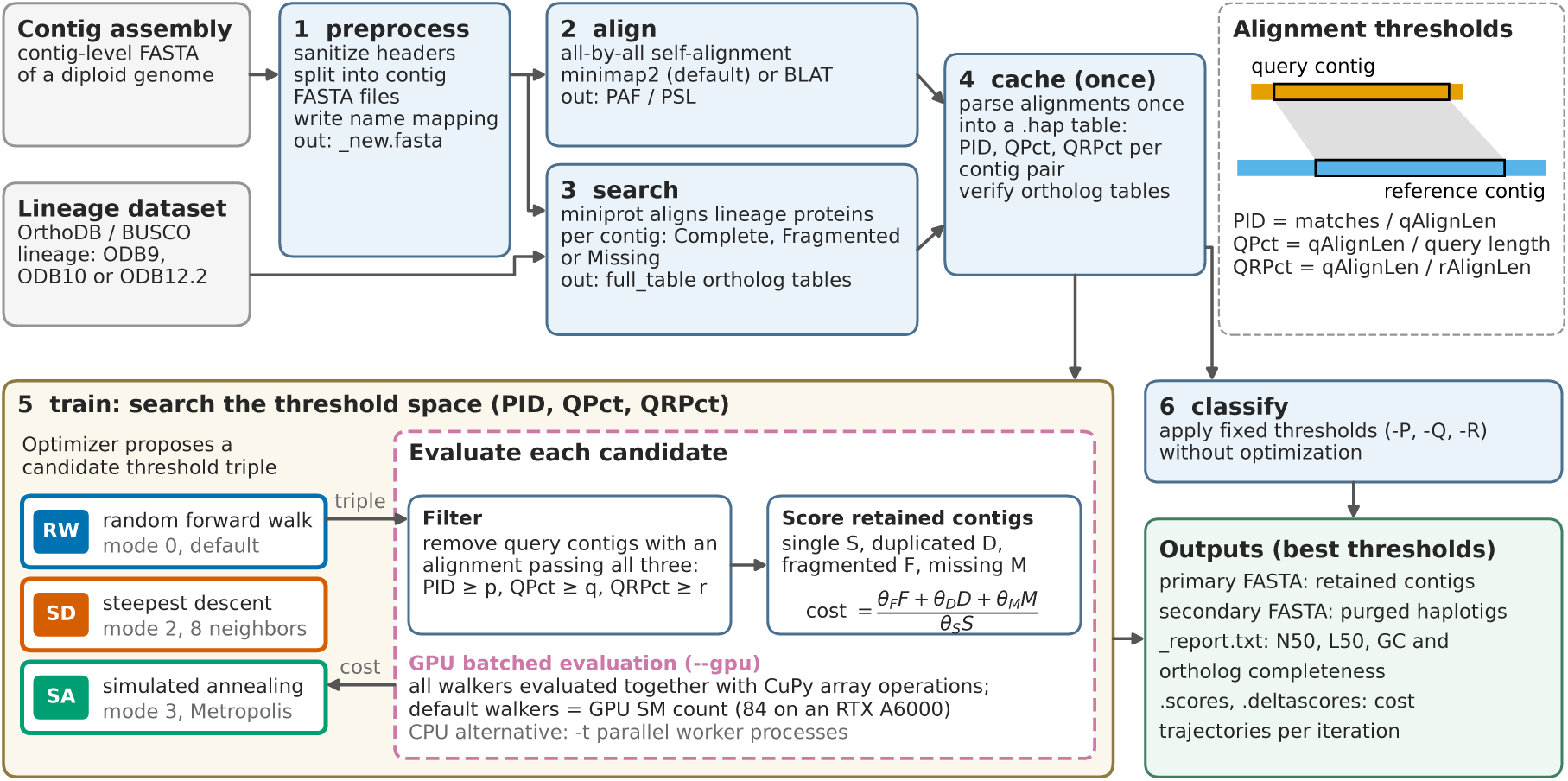
Workflow of HapSolo2, updated from Fig. 1 of Solares et al. (2021). The contig assembly is preprocessed (1), aligned to itself (2) and searched for the orthologs of a lineage dataset (3); the alignments are parsed once into a cache of PID, QPct and QRPct per contig pair (4). Training (5) searches the threshold space with RW, SD or SA: for each candidate threshold triple, every query contig with an alignment that passes all three thresholds is removed, and the retained contigs are scored by their single-copy (S), duplicated (D), fragmented (F) and missing (M) orthologs. With `--gpu`, all walkers are evaluated together on the GPU. Classification (6) applies fixed thresholds without optimization. The inset defines the three alignment metrics. Both steps write the primary and secondary assemblies, a statistics report, and the cost trajectory of every walker.

### Pipeline and command-line interface

HapSolo2 is driven by one command, `hapsolo-cli`, with six subcommands that follow the workflow of Figure 1. `preprocess` sanitizes FASTA headers (the first whitespace-delimited token is kept, characters outside `[a-zA-Z0-9.]` are replaced by underscores, and duplicate names are resolved by truncation to the shortest unique prefix), writes the sanitized assembly and per-contig FASTA files for contigs below a size limit (10 Mb by default), and records the mapping from original to sanitized names. `align` runs the all-by-all self-alignment with minimap2 (default) or BLAT and compresses the output. `search` aligns the ortholog proteins of a lineage dataset to the assembly with miniprot and classifies each ortholog on each contig. `cache` parses the alignment file once into a compact `.hap` table and verifies the ortholog tables, so that repeated training runs skip the parsing of multi-gigabyte PAF files. `train` runs the optimizer and writes the primary and secondary assemblies, a statistics report, and the cost trajectory of every walker. `classify` applies a fixed threshold triple without optimization, to reproduce a previous result or to apply trained thresholds to a new assembly. Each step is also available as a Python module, and the root-level scripts of the first release remain as backward-compatible stubs. Contig names are reconciled across the FASTA, alignment and ortholog files by exact match, by match after sanitization, and by a unique prefix relation, so that users who skip preprocessing are not blocked by header differences.

### Ortholog search and classification

The original HapSolo ran BUSCO v3.0.2 in genome mode on each contig separately. HapSolo2 instead aligns the ortholog proteins of a BUSCO lineage dataset to the whole assembly in a single run of miniprot (Li 2023) in PAF mode, with the maximum intron length set from the genome length (option `-I`, 3.6 times the square root of the genome length, bounded between 10 kb and 300 kb). The datasets are the per-lineage subsets of OrthoDB curated by the BUSCO team (Kriventseva et al. 2019; Manni et al. 2021; Tegenfeldt et al. 2025), and HapSolo2 reads the three formats in use: ODB9 (`ancestral_variants` or `ancestral` protein sets) and ODB10 and ODB12.2 (`refseq_db.faa`, gzipped or not). For ODB10 and later, protein identifiers are mapped to ortholog groups by trying underscore-delimited prefixes of the identifier, shortest first, against the keys of the dataset’s `scores_cutoff` file; proteins whose group has no score cutoff are excluded with a warning. For each ortholog group and contig, the highest-scoring alignment is retained. An alignment is significant if its miniprot alignment score meets the group’s score cutoff; a significant alignment is Complete if the aligned protein length reaches the group’s expected length from `lengths_cutoff`, two scale units below the group’s central length (the mean minus two standard deviations in ODB9 and ODB10 datasets, and the median minus two scaled median absolute deviations in ODB12.2 datasets; BUSCO development team 2026), and Fragmented otherwise; a group with no significant alignment on any contig is Missing. The released classifier applies three further filters, set with `--classify-params`: a minimum ratio of score to cutoff (2.5), a minimum alignment coverage (0.7), and a minimum score of a second contig relative to the best contig (0.9) for the group to count on that contig as well. The ortholog tables used for the reported results were produced by the July 2026 classifier, which did not apply these filters (Methods). The per-contig results are written in the BUSCO `full_table` format, so that the optimizer accepts either HapSolo2 classifications or BUSCO v3 output from the original pipeline.

### Alignment metrics and the removal rule

For every alignment record with aligned query span *L_q_* and aligned reference span *L_r_*, PID is the number of residue matches divided by *L_q_*, QPct is *L_q_* divided by the query length, and QRPct is *L_q_* divided by *L_r_*. PID is therefore normalized by the aligned query span rather than by the alignment block length. Records are discarded if they are self-alignments, if the query contig is shorter than 1,000 bp, or if any of the three metrics falls below 0.2; the retained records (query, target, query length, QPct, PID, QRPct) are written to the `.hap` cache, and the prefilter thresholds define the lower bounds of the search space. For a threshold triple (*p, q, r*), a record hits if

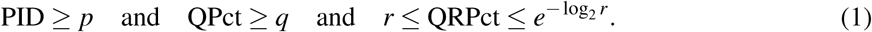

The upper bound *e*^−log2^ *^r^* = *r*^−1^*^/^*^ln2^ ≈ *r*^−1^.^443^ is the symmetric counterpart QR^′^ of the original description; for *r* = 0.5 it equals *e* ≈ 2.718, so the window admits alignments whose query and reference spans are of similar length and narrows as *r* approaches 1. The query contig of every hitting record is removed. The self-alignment reports both orientations of an overlapping pair, so both contigs of a pair are removed when both directed records hit. Contigs that never appear as a query among the retained records are always kept, contigs shorter than 1,000 bp (`--min-contig`) are always excluded from the primary assembly, and the removed contigs form the secondary assembly.

### Cost function

Ortholog membership is encoded once at startup as an integer matrix of ortholog groups by contigs, with 0 for absent, 1 for Complete and 2 for Fragmented. For a kept set of contigs, a group is single-copy (S) if it is Complete on exactly one kept contig, duplicated (D) if Complete on more than one, fragmented (F) if Complete on none and Fragmented on at least one, and missing (M) otherwise; a group that is Complete on more than one retained contig is the signature of a retained haplotig. The cost of a candidate is

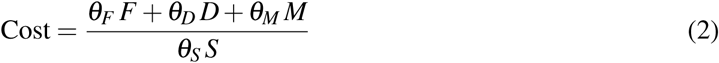

with default weights θ*_S_* = θ*_D_*= θ*_M_*= 1 and θ*_F_* = 0, as in the original. Lower is better: the cost rewards single-copy orthologs and penalizes both retained haplotigs and removed primary sequence. A candidate with *S* = 0 receives a cost of 5 × 10^7^. The weights are set with `-S`, `-D`, `-F` and `-M`, and an arbitrary expression in S, D, F, M, C and *n* (the number of groups) can be supplied with `--formula`; the expression is parsed against a whitelist of variables, operators and functions before evaluation. All benchmark runs in this paper used the default weights. We recommend `-M 2 -D 1.5` as a starting point, which penalizes missing orthologs twice and duplicated orthologs 1.5 times relative to the defaults and encourages the optimizer to retain contigs that carry unique orthologs while still removing redundant haplotigs.

### Optimizers

The search space is *p, q, r* ∈ [0.2, 1]. The step size *s* and the plateau resolution are both 1 × 10^−4^. Every walker starts at a point drawn uniformly from the search space, is on a plateau when the cost has changed by no more than the resolution over the preceding nine iterations, and redraws a coordinate uniformly when it leaves the space above 1.0. The iteration budget is given per walker (`-n`) or in total (`--totaliters`), in which case it is divided evenly among walkers. Three optimizers share this framework and are selected with `--mode`.

#### Random forward walk (mode 0, RW)

Each coordinate is incremented by 0, *s* or 2*s*, drawn independently and uniformly with the all-zero step redrawn, and the walker always moves to the proposal. The walk is monotone in all three thresholds, so each walker traces a path from permissive to stringent values, and the search relies on many walkers and on restarts, at a plateau or when all three coordinates exceed 1, to cover the space. This is the strategy of the original HapSolo, reimplemented over the vectorized evaluation core.

#### Steepest descent (mode 2, SD)

At each iteration all eight diagonal neighbors (*p* ± *s*, *q* ± *s*, *r* ± *s*) inside the search space are evaluated and the walker moves to the lowest-cost neighbor, including when every neighbor is worse than the current point, so the method is a greedy best-neighbor walk. Restarts occur at plateaus and when no neighbor is in bounds. Each iteration costs up to eight evaluations.

#### Simulated annealing (mode 3, SA)

Proposals use the forward steps of RW. A proposal that does not increase the cost is always accepted; a proposal that increases the cost by δ is accepted with probability exp(−δ */T*) (Metropolis criterion; Kirkpatrick et al. 1983), and a rejected proposal leaves the walker in place. Unless set with `--sa-temp`, the initial temperature is calibrated from the data: ten points are drawn uniformly, σ is the standard deviation of the absolute cost differences between consecutive points, and

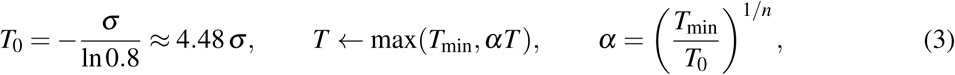

so that a worsening of size σ is accepted with probability 0.8 at the start, and the geometric schedule reaches *T*_min_ = 1 ×10^−6^ at the last of the *n* iterations per walker (`--sa-cooling`, `--sa-min-temp`). A plateau triggers a restart from a uniform point and a reheat. In the code used for the reported results (July 2026), *T* is reset to 0.5 *T*_0_ at every plateau and cooled once in the same step. In the released code, *T* is reset to *T*_0_ × 0.5*^k^*^−1^ at the *k*-th plateau of a walker, bounded below by *T*_min_, without cooling in that step.

#### Fixed thresholds (mode 1)

A single candidate is evaluated at user-supplied thresholds and written out; this mode underlies `hapsolo-cli classify`.

### Retention of candidate assemblies

Every evaluated candidate is offered to a best-*B* queue that keeps the *B* lowest-cost candidates with distinct kept sets (*B* = 1 by default, option `-B`), the unique priority queue of the original HapSolo. The queue is seeded with the unfiltered assembly, so that no run can return a result worse than no reduction. The queue stays on the host in all modes: it holds variable-size contig sets, is consulted once per iteration, and is not a computational bottleneck.

### GPU implementation

Profiling of the original implementation showed that the cost of an iteration is dominated by re-deriving the kept set and recounting orthologs, not by the threshold comparison itself. The GPU implementation therefore keeps all alignment and ortholog data resident on the device and evaluates the candidates of every walker in a single batched operation per iteration, so that the number of kernel launches per iteration is independent of the number of walkers. It uses CuPy (Okuta et al. 2017) without cuDF or hand-written kernels, and runs in one process and one CUDA context on one device.

#### Data residency

The following are transferred to the device once per run: the PID, QPct and QRPct columns of the alignment cache as float32 vectors of length *A* (the number of alignment records); the query-contig index of each record (int32); boolean masks over the *C* contigs for all contigs, for contigs that are never a query, and for contigs shorter than 1,000 bp; and the ortholog matrix (uint8, *O* ×*C* for *O* groups).

#### Batched evaluation

For *N* candidates with thresholds held as length-*N* vectors, each iteration (1) computes the hit matrix *H* (*A* × *N*, boolean) by broadcasting the four comparisons of the removal rule over all records and candidates; (2) scatters hits to a removal mask (*N* × *C*) through the query-contig index of each hitting record; (3) forms the kept mask (*N* × *C*) as not removed or never a query, excluding short contigs; (4) masks the ortholog matrix with the kept mask into an *N* × *O* × *C* uint8 tensor and reduces over contigs to obtain Complete and Fragmented counts per ortholog, from which S, D, F and M follow; and (5) evaluates the cost for all *N* candidates. The *N* × *O* × *C* tensor is the memory bottleneck, so candidates are processed in chunks sized to the free device memory, chunk = max 16, ⌊0.6 × free memory*/*(*O* ×*C* × 5 bytes)⌋, and on an out-of-memory error the chunk is halved and retried up to three times. In this path the walker logic (proposals, plateau tests, acceptance, restarts) runs on the host, and the *N* costs are copied to the host once per iteration. SD submits up to 8*N* candidates per iteration and so gives the GPU eight times more work per launch than RW or SA.

#### Synchronization-free loop

The released code adds a second path for RW and SA that keeps the entire walker loop on the device. Three changes remove host-device synchronization from the loop. First, the removal mask is computed as a sparse-dense product *R* = *SH*, where *S* is a *C* × *A* compressed sparse row matrix with one nonzero per column at the record’s query contig, so that *R*[*c, n*] counts the hits on contig *c* for candidate *n* and removal is *R >* 0; this replaces a nonzero scan that may synchronize the device. Second, Complete and Fragmented indicator matrices (*O* × *C*, float32) are multiplied by the transposed kept mask, giving per-ortholog counts for all candidates in two matrix products without forming the *N* × *O* ×*C* tensor; the inputs are 0 or 1 and the counts are at most *C*, so the float32 results are exact integers for *C <* 2^24^. Third, proposals, bounds handling, Metropolis acceptance and best-so-far tracking are expressed as element-wise selects on device arrays, random numbers are drawn with CuPy’s generator, and the host reads the device only to refresh the progress display every 200 iterations. This path applies the same plateau restart, SA reheat and forward-only RW steps as the CPU optimizers and returns the cost history of every walker. The *A. radiata* runs on the complete alignment cache used this path for RW and SA and the batched path for SD; the runs for the other three species used the batched path of the July 2026 code.

#### Number of walkers

The default number of walkers equals the number of streaming multiprocessors of the device (84 on an RTX A6000, 108 on an A100), so that every multiprocessor receives work in each batched kernel. A user-supplied `--agents` is capped at ⌊0.6 ×device memory*/*(*O* ×*C* × 5 bytes)⌋, the size of the dense scoring tensor, and the cap is reported; the default is not capped.

### CPU implementation

The CPU path evaluates candidates with the same rule in numpy: the alignment columns are float32 arrays, hits are found with a vectorized four-way comparison, and ortholog counts are computed from the integer matrix. Walkers run in a `multiprocessing` pool with the fork start method, inheriting the read-only data from the parent process. Neither path uses pandas in the optimization loop, and threshold comparisons are made in float32 on both, so a CPU run and a GPU run evaluate the same candidate identically.

### Output, assembly statistics, and plotting

`train` and `classify` write the primary and secondary assemblies with contigs in the order of the input assembly, so that identical runs give byte-identical files; a report with contig counts at several size thresholds, total length, largest contig, N50, N75, L50, L75, GC content, N density and ortholog completeness; and `.scores` and `.deltascores` files holding the cost trajectory of every walker, for convergence plots. The selected threshold values are encoded in the output file names. A separate tool, `assembly_stats.py`, reports the same contiguity statistics for one or more FASTA files side by side, adds NG50, NG75, LG50 and LG75 for a user-supplied genome size and the size and genome fraction of the largest *n* sequences for a user-supplied chromosome number, and draws a cumulative length plot (PNG and PDF), for example to compare contigs with the Hi-C scaffolds built from them. Together the report and the tool replace the separate QUAST (Gurevich et al. 2013) and BUSCO (Simão et al. 2015) runs of the original post-processing.

### Distribution and requirements

HapSolo2 is written for Python 3 and depends on numpy, pandas and tqdm, with CuPy as an optional dependency for GPU mode; `pip install.` installs the `hapsolo` and `hapsolo-cli` entry points and `pip install .[gpu]` adds CuPy. minimap2 (Li 2018) (or BLAT; Kent 2002) and miniprot must be on the path. An Apptainer definition file (`hapsolo2.def`) builds an image that bundles HapSolo2, minimap2 2.31, miniprot 0.18, BLAT, `assembly_stats.py` and the CUDA 12 runtime, so that only the NVIDIA driver is needed on the host; the image ignores packages installed in the user’s home directory and records the versions it carries. ODB10 and ODB12.2 lineage datasets are downloaded from the BUSCO data archive. The ODB9 datasets, which the BUSCO team no longer distributes, we have archived on Zenodo with checksums (Zdobnov et al. 2026, https://doi.org/10.5281/zenodo.20987183), so that results produced with the original HapSolo remain reproducible. ODB12 (OrthoDB 12.1) datasets are read but not recommended, since they lack length cutoffs and their group identifiers are not retained in ODB12.2 (BUSCO development team 2026). SLURM batch scripts for CPU and GPU runs are included, and the package carries 435 tests across the optimizer modes, the search step, the command-line interface, and real-data integration. HapSolo2 is released under the Apache License 2.0 at https://github.com/esolares/HapSolo.

## Results

### Input assemblies

The four assemblies span a 15-fold range in size and a 24-fold range in contig number (Table 1), from 212.0 Mb in 1,073 contigs (*A. funestus*) to 3,229.4 Mb in 16,218 contigs (*A. radiata*). Their duplication differs even more. BUSCO v6.1 against the odb12.2 lineages classes 52.0% of the *P. americana* orthologs as duplicated, consistent with both haplotypes of most of the genome being retained in the primary assembly, against 23.2% for *A. radiata*, 17.9% for *V. vinifera* and 7.8% for *A. funestus*. Completeness is high in all four (92.0% to 99.6%), so the duplicated fraction, not missing genes, is what HapSolo2 has to correct. The *A. funestus* and *A. radiata* assemblies are the files analyzed in Solares et al. (2021). The *V. vinifera* assembly is a 684-contig Chardonnay assembly rather than the 2,072-contig hybrid assembly analyzed with HapSolo, and the *P. americana* cv. Gwen contig assembly is that of Solares et al. (2023) before haplotig removal.

**Table 1:** Input assemblies and independent BUSCO assessment (v6.1, genome mode, odb12.2).

| Species | Contigs | Size (Mb) | Contig N50 (kb) | BUSCO lineage | C (%) | S (%) | D (%) | F (%) | M (%) |
| --- | --- | --- | --- | --- | --- | --- | --- | --- | --- |
| *Anopheles funestus* | 1,073 | 212.0 | 630 | diptera_odb12.2 | 95.9 | 88.1 | 7.8 | 0.2 | 3.9 |
| *Persea americana* cv. Gwen | 5,122 | 1,449.9 | 1,428 | embryophyta_odb12.2 | 99.6 | 47.6 | 52.0 | 0.0 | 0.4 |
| *Vitis vinifera* cv. Chardonnay | 684 | 606.0 | 24,528 | embryophyta_odb12.2 | 98.6 | 80.7 | 17.9 | 0.2 | 1.3 |
| *Amblyraja radiata* | 16,218 | 3,229.4 | 427 | vertebrata_odb12.2 | 92.0 | 68.9 | 23.2 | 2.8 | 5.2 |

### Ortholog datasets

Each assembly was trained on three OrthoDB releases (ODB9, ODB10 and ODB12.2) of one lineage, diptera for *A. funestus* and embryophyta for the two plants, and *A. radiata* was trained on two lineages (actinopterygii and metazoa), since no dedicated lineage exists for cartilaginous fishes. The datasets differ in size (932 to 9,048 groups) and in classification behavior. ODB10 classified no ortholog as Fragmented in any of the 15 ODB10 runs (Table 2), so its internal C, D and M are inflated relative to the other releases. ODB12 (OrthoDB 12.1) was evaluated and excluded: lacking length cutoffs, it classified 94.1% of the *P. americana* orthologs as Fragmented and produced a misleadingly low cost (0.053) on a near-meaningless classification. Internal costs are therefore compared only within a training release, and assemblies trained on different releases are compared with an independent BUSCO assessment on a common lineage (below).

**Table 2:** HapSolo2 GPU runs (100,000 iterations): internal ortholog scores on the training release. Cost = (*D* + *M*)*/S*. C, S, D, F and M are HapSolo2’s miniprot-based counts on the training release as a percentage of *n* groups; costs are comparable only within one training release. Wall = end-to-end wall time in seconds (RW and SD from the timed run logs; SA from the SA sweep; † = value from the results table rather than a timed log).

| ODB ($n$) | Opt. | Cost | PID | QPct | QRPct | Contigs | Size (Mb) | Removed (%) | N50 (kb) | C (%) | S (%) | D (%) | F (%) | M (%) | Wall (s) |
| --- | --- | --- | --- | --- | --- | --- | --- | --- | --- | --- | --- | --- | --- | --- | --- |
| ODB9 (2,799) | RW | 0.0519 | 0.5969 | 0.7896 | 0.7524 | 906 | 205.4 | 3.1 | 669 | 95.2 | 93.5 | 1.7 | 1.6 | 3.2 | 23 |
| ODB9 (2,799) | SD | 0.0528 | 0.5860 | 0.7996 | 0.6544 | 903 | 205.3 | 3.2 | 669 | 95.1 | 93.5 | 1.6 | 1.6 | 3.3 | 90 |
| ODB9 (2,799) | SA | 0.0524 | 0.5829 | 0.7921 | 0.7430 | 904 | 205.3 | 3.2 | 669 | 95.2 | 93.5 | 1.7 | 1.6 | 3.2 | 24 |
| ODB10 (3,285) | RW | 0.1087 | 0.2958 | 0.8536 | 0.7607 | 888 | 205.1 | 3.2 | 669 | 97.7 | 90.2 | 7.5 | 0.0 | 2.3 | 25 |
| ODB10 (3,285) | SD | 0.1087 | 0.3388 | 0.8571 | 0.7412 | 893 | 205.3 | 3.2 | 669 | 97.7 | 90.2 | 7.5 | 0.0 | 2.3 | 112 |
| ODB10 (3,285) | SA | 0.1087 | 0.2002 | 0.8536 | 0.7607 | 879 | 204.9 | 3.3 | 669 | 97.7 | 90.2 | 7.5 | 0.0 | 2.3 | 25 |
| ODB12.2 (3,914) | RW | 0.0996 | 0.2755 | 0.7967 | 0.4012 | 854 | 204.0 | 3.8 | 687 | 94.5 | 87.7 | 6.8 | 3.5 | 2.0 | 26 |
| ODB12.2 (3,914) | SD | 0.0996 | 0.3306 | 0.7997 | 0.4405 | 864 | 204.2 | 3.7 | 669 | 94.5 | 87.7 | 6.8 | 3.5 | 2.0 | 129 |
| ODB12.2 (3,914) | SA | 0.0996 | 0.3143 | 0.8015 | 0.2029 | 861 | 204.1 | 3.7 | 669 | 94.5 | 87.7 | 6.8 | 3.5 | 2.0 | 27 |

**(b) *Persea americana* cv. Gwen, embryophyta training lineage**
| ODB ($n$) | Opt. | Cost | PID | QPct | QRPct | Contigs | Size (Mb) | Removed (%) | N50 (kb) | C (%) | S (%) | D (%) | F (%) | M (%) | Wall (s) |
| --- | --- | --- | --- | --- | --- | --- | --- | --- | --- | --- | --- | --- | --- | --- | --- |
| ODB9 (1,440) | RW | 0.1036 | 0.2214 | 0.3660 | 0.2792 | 496 | 860.5 | 40.7 | 4,608 | 95.2 | 89.8 | 5.4 | 0.9 | 3.9 | 66 |
| ODB9 (1,440) | SD | 0.1036 | 0.2557 | 0.3663 | 0.2246 | 514 | 862.5 | 40.5 | 4,608 | 95.2 | 89.8 | 5.4 | 0.9 | 3.9 | 225 |
| ODB9 (1,440) | SA | 0.1045 | 0.4355 | 0.3657 | 0.5177 | 618 | 871.7 | 39.9 | 4,601 | 95.2 | 89.7 | 5.5 | 0.9 | 3.9 | 61 |
| ODB10 (1,614) | RW | 0.3350 | 0.2683 | 0.3638 | 0.6286 | 518 | 861.2 | 40.6 | 4,608 | 98.7 | 74.9 | 23.8 | 0.0 | 1.3 | 61 |
| ODB10 (1,614) | SD | 0.3350 | 0.2110 | 0.3642 | 0.2424 | 490 | 859.3 | 40.7 | 4,608 | 98.7 | 74.9 | 23.8 | 0.0 | 1.3 | 255 |
| ODB10 (1,614) | SA | 0.3350 | 0.2181 | 0.3590 | 0.6474 | 487 | 857.4 | 40.9 | 4,608 | 98.7 | 74.9 | 23.8 | 0.0 | 1.3 | 59 |
| ODB12.2 (1,658) | RW | 0.4372 | 0.2811 | 0.3435 | 0.3180 | 493 | 849.9 | 41.4 | 4,636 | 98.6 | 69.1 | 29.5 | 0.7 | 0.7 | 64 |
| ODB12.2 (1,658) | SD | 0.4372 | 0.3431 | 0.3443 | 0.6321 | 523 | 852.6 | 41.2 | 4,636 | 98.6 | 69.1 | 29.5 | 0.7 | 0.7 | 246 |
| ODB12.2 (1,658) | SA | 0.4372 | 0.2510 | 0.3435 | 0.3013 | 477 | 848.5 | 41.5 | 4,636 | 98.6 | 69.1 | 29.5 | 0.7 | 0.7 | 62 |

**(c) *Vitis vinifera* cv. Chardonnay, embryophyta training lineage**
| ODB (*n*) | Opt. | Cost | PID | QPct | QRPct | Contigs | Size (Mb) | Removed (%) | N50 (kb) | C (%) | S (%) | D (%) | F (%) | M (%) | Wall (s) |
| --- | --- | --- | --- | --- | --- | --- | --- | --- | --- | --- | --- | --- | --- | --- | --- |
| ODB9 (1,440) | RW | 0.0386 | 0.5822 | 0.2251 | 0.6706 | 112 | 517.9 | 14.5 | 25,768 | 97.4 | 95.4 | 2.0 | 0.9 | 1.7 | 153† |
| ODB9 (1,440) | SD | 0.0386 | 0.5843 | 0.2290 | 0.2419 | 112 | 518.7 | 14.4 | 25,768 | 97.4 | 95.4 | 2.0 | 0.9 | 1.7 | 184† |
| ODB9 (1,440) | SA | 0.0393 | 0.4551 | 0.2081 | 0.4461 | 67 | 511.2 | 15.6 | 25,768 | 97.4 | 95.3 | 2.0 | 0.9 | 1.7 | 26 |
| ODB10 (1,614) | RW | 0.2227 | 0.2459 | 0.2246 | 0.4845 | 53 | 509.7 | 15.9 | 25,768 | 99.3 | 81.8 | 17.5 | 0.0 | 0.7 | 138† |
| ODB10 (1,614) | SD | 0.2227 | 0.3119 | 0.2103 | 0.5265 | 54 | 509.2 | 16.0 | 25,768 | 99.3 | 81.8 | 17.5 | 0.0 | 0.7 | 178† |
| ODB10 (1,614) | SA | 0.2227 | 0.2244 | 0.2227 | 0.2896 | 53 | 509.7 | 15.9 | 25,768 | 99.3 | 81.8 | 17.5 | 0.0 | 0.7 | 28 |
| ODB12.2 (1,658) | RW | 0.2654 | 0.2860 | 0.2156 | 0.4498 | 56 | 509.5 | 15.9 | 25,768 | 98.7 | 78.2 | 20.5 | 1.1 | 0.2 | 141† |
| ODB12.2 (1,658) | SD | 0.2654 | 0.2644 | 0.2213 | 0.6604 | 52 | 509.3 | 16.0 | 25,768 | 98.7 | 78.2 | 20.5 | 1.1 | 0.2 | 181† |
| ODB12.2 (1,658) | SA | 0.2654 | 0.2322 | 0.2072 | 0.5201 | 47 | 508.6 | 16.1 | 25,768 | 98.7 | 78.2 | 20.5 | 1.1 | 0.2 | 31 |

**(d) *Amblyraja radiata*, actinopterygii training lineage**
| ODB (*n*) | Opt. | Cost | PID | QPct | QRPct | Contigs | Size (Mb) | Removed (%) | N50 (kb) | C (%) | S (%) | D (%) | F (%) | M (%) | Wall (s) |
| --- | --- | --- | --- | --- | --- | --- | --- | --- | --- | --- | --- | --- | --- | --- | --- |
| ODB9 (4,584) | RW | 0.3029 | 0.7825 | 0.7856 | 0.8832 | 16,138 | 3,186.6 | 1.3 | 421 | 86.3 | 72.4 | 13.9 | 5.6 | 8.1 | 292 |
| ODB9 (4,584) | SD | 0.3025 | 0.7826 | 0.7659 | 0.9323 | 16,135 | 3,184.0 | 1.4 | 420 | 86.3 | 72.5 | 13.9 | 5.6 | 8.1 | 1,696 |
| ODB9 (4,584) | SA | 0.3029 | 0.7852 | 0.7877 | 0.4365 | 16,140 | 3,188.0 | 1.3 | 421 | 86.3 | 72.4 | 13.9 | 5.6 | 8.1 | 320 |
| ODB10 (3,640) | RW | 0.4543 | 0.2476 | 0.7155 | 0.9486 | 16,025 | 3,117.1 | 3.5 | 407 | 89.0 | 68.8 | 20.3 | 0.0 | 11.0 | 280 |
| ODB10 (3,640) | SD | 0.4543 | 0.6670 | 0.7154 | 0.5340 | 16,026 | 3,118.9 | 3.4 | 408 | 89.0 | 68.8 | 20.3 | 0.0 | 11.0 | 1,373 |
| ODB10 (3,640) | SA | 0.4548 | 0.2781 | 0.7280 | 0.9572 | 16,037 | 3,123.4 | 3.3 | 409 | 89.1 | 68.7 | 20.3 | 0.0 | 10.9 | 287 |
| ODB12.2 (9,048) | RW | 0.4575 | 0.6765 | 0.5988 | 0.7536 | 15,963 | 3,079.7 | 4.6 | 400 | 73.2 | 54.7 | 18.5 | 20.3 | 6.5 | 493 |
| ODB12.2 (9,048) | SD | 0.4582 | 0.6605 | 0.5985 | 0.4120 | 15,957 | 3,076.4 | 4.7 | 399 | 73.2 | 54.7 | 18.5 | 20.3 | 6.5 | 3,206 |
| ODB12.2 (9,048) | SA | 0.4573 | 0.6844 | 0.5058 | 0.3576 | 15,884 | 3,026.7 | 6.3 | 390 | 72.4 | 54.6 | 17.8 | 20.4 | 7.2 | 530 |

(e) *Amblyraja radiata*, metazoa training lineage
| ODB (*n*) | Opt. | Cost | PID | QPct | QRPct | Contigs | Size (Mb) | Removed (%) | N50 (kb) | C (%) | S (%) | D (%) | F (%) | M (%) | Wall (s) |
| --- | --- | --- | --- | --- | --- | --- | --- | --- | --- | --- | --- | --- | --- | --- | --- |
| ODB9 (978) | RW | 0.3138 | 0.6828 | 0.4401 | 0.2297 | 15,800 | 2,966.7 | 8.1 | 379 | 84.3 | 71.4 | 12.9 | 6.2 | 9.5 | 157 |
| ODB9 (978) | SD | 0.3138 | 0.6308 | 0.4384 | 0.8891 | 15,776 | 2,951.0 | 8.6 | 376 | 84.3 | 71.4 | 12.9 | 6.2 | 9.5 | 421 |
| ODB9 (978) | SA | 0.3138 | 0.6273 | 0.4396 | 0.6533 | 15,771 | 2,948.2 | 8.7 | 375 | 84.3 | 71.4 | 12.9 | 6.2 | 9.5 | 157 |
| ODB10 (954) | RW | 2.9750 | 0.4380 | 0.2267 | 0.8980 | 15,229 | 2,535.8 | 21.5 | 307 | 95.0 | 25.2 | 69.8 | 0.0 | 5.0 | 131 |
| ODB10 (954) | SD | 2.9259 | 0.6264 | 0.2276 | 0.9360 | 15,270 | 2,559.5 | 20.7 | 310 | 95.4 | 25.5 | 69.9 | 0.0 | 4.6 | 423 |
| ODB10 (954) | SA | 2.9750 | 0.2894 | 0.2267 | 0.9031 | 15,228 | 2,535.2 | 21.5 | 306 | 95.0 | 25.2 | 69.8 | 0.0 | 5.0 | 151 |
| ODB12.2 (932) | RW | 1.0152 | 0.6477 | 0.2043 | 0.8144 | 15,138 | 2,462.9 | 23.7 | 295 | 83.0 | 42.3 | 40.8 | 14.8 | 2.1 | 130 |
| ODB12.2 (932) | SD | 1.0102 | 0.4426 | 0.2042 | 0.9106 | 15,114 | 2,446.1 | 24.3 | 293 | 82.7 | 42.2 | 40.6 | 15.2 | 2.0 | 429 |
| ODB12.2 (932) | SA | 1.0152 | 0.6481 | 0.2043 | 0.3571 | 15,139 | 2,463.4 | 23.7 | 295 | 83.0 | 42.3 | 40.8 | 14.8 | 2.1 | 132 |

### Optimization results

Table 2 lists the 45 GPU runs of 100,000 iterations each. Two points stand out. First, the three optimizers reach nearly the same optimum. Within a species, lineage and release, the best costs of RW, SD and SA differ by at most 1.8% in all 15 combinations and are identical to four decimals in 7. Cost differences of this size correspond to a handful of orthologs; for *A. funestus* with ODB9, for example, the RW optimum (0.0519) and the SA optimum (0.0524) both retain 93.5% single-copy orthologs. SD reached a strictly lower cost than both RW and SA in 3 combinations, all on *A. radiata* (actinopterygii ODB9, metazoa ODB10 and metazoa ODB12.2), by at most 1.7%. The threshold triples that reach these optima, however, differ widely between optimizers (PID of 0.2214, 0.2557 and 0.4355 for the three *P. americana* ODB9 runs, for example), which indicates that the cost landscape is dominated by broad basins in which many threshold triples retain the same ortholog-bearing contigs.

Second, the amount removed reflects the duplication of the assembly. HapSolo2 removed 39.9% to 41.5% of the *P. americana* assembly (from 1,449.9 Mb to 848.5 to 871.7 Mb, and from 5,122 to 477 to 618 contigs) and raised contig N50 from 1,428 kb to 4,601 to 4,636 kb, a 3.2-fold increase. It removed 14.4% to 16.1% of *V. vinifera* (606.0 Mb to 508.6 to 518.7 Mb; 684 to 47 to 112 contigs) and 3.1% to 3.8% of *A. funestus* (212.0 Mb to 204.0 to 205.4 Mb). For *A. radiata*, removal depended strongly on the training lineage: 1.3% to 6.3% with actinopterygii against 8.1% to 24.3% with metazoa. With metazoa ODB10 and ODB12.2, the internal duplicated fraction remained at 40.6% to 69.9% while 20.7% to 24.3% of the assembly was removed; these runs have the highest costs in Table 2 (1.01 to 2.98), and their independent assessment is pending (below). Figure 2 plots the cost trajectories of the three optimizers and Figure 3 the cumulative contig length distributions of the unreduced and reduced assemblies.

**Figure 2:**
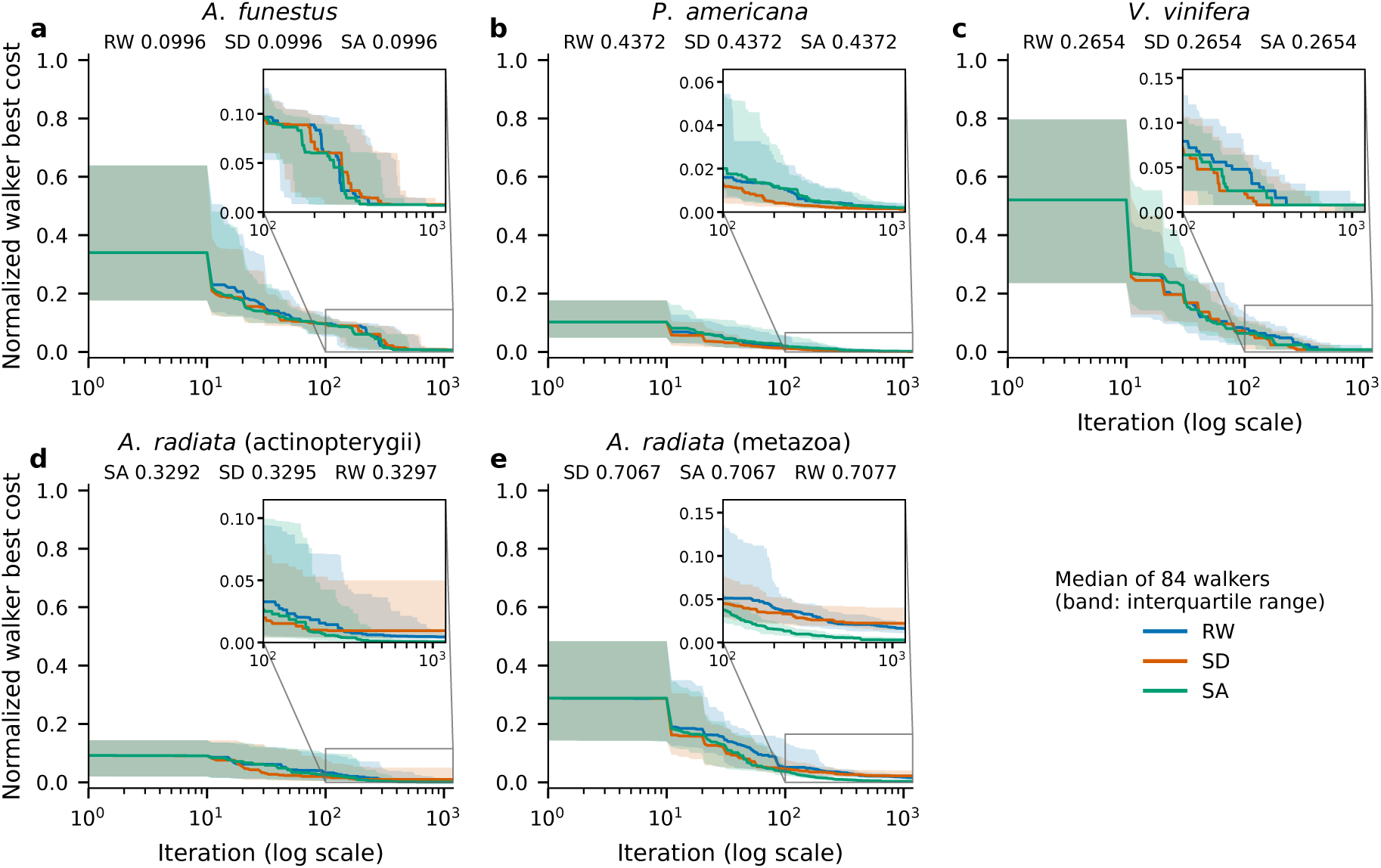
Convergence of RW, SD and SA on each species and training lineage with ODB12.2 (GPU, 84 walkers × 1,191 iterations, 100,000 total iterations), from the `.scores` files. For each walker the best cost reached so far is tracked at every iteration; lines show the median across the 84 walkers and shaded bands the interquartile range. Costs are normalized per panel as (Cost - *C*_min_)*/*(*C*_max_ - *C*_min_), where *C*_min_ is the lowest cost reached by any of the three optimizers and *C*_max_ the highest starting cost, so that the three optimizers, which share the same 84 start points, are drawn on one scale, as in Solares et al. (2021). Insets enlarge iterations 100 to 1,191. The minimum raw cost of each optimizer is printed under each panel title, lowest first. (a) *A. funestus*, diptera. (b) *P. americana*, embryophyta. (c) *V. vinifera*, embryophyta. (d) *A. radiata*, actinopterygii. (e) *A. radiata*, metazoa. Supplementary Figure S1 shows ODB9 and ODB10.

**Figure 3:**
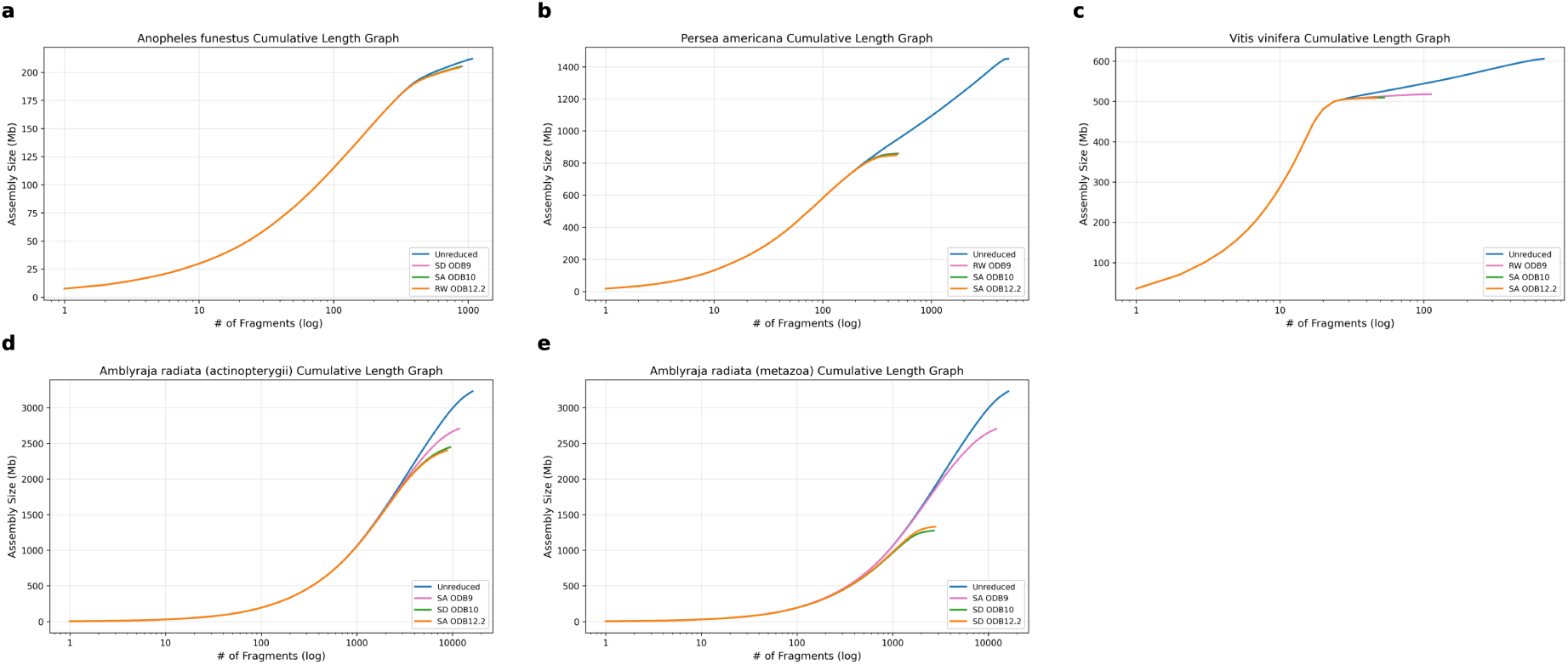
Cumulative contig length of the unreduced assembly and of the best HapSolo2 assembly for each training release (ODB9, ODB10 and ODB12.2), drawn with `assembly_stats.py`. Contigs are sorted from longest to shortest, and the cumulative assembly size (Mb) is plotted against the number of contigs (log scale); the legend names the optimizer of each selected run. (a) *A. funestus*, diptera. (b) *P. americana*, embryophyta. (c) *V. vinifera*, embryophyta. (d) *A. radiata*, actinopterygii. (e) *A. radiata*, metazoa. *A. radiata* runs are selected by internal cost on the complete alignment cache.

### Independent validation with BUSCO

HapSolo2’s internal scores are computed on the training release and are not comparable across releases, so the reduced assemblies were assessed with BUSCO v6.1 in genome mode against one odb12.2 lineage per species (Table 3). The assessment is independent of the training release in its lineage dataset and its classification rules, although BUSCO’s genome mode maps proteins with miniprot, as HapSolo2 does. For each species, lineage and release, the run with the highest *C* + *S* − 2*D* on the internal scores was assessed. In every species assessed so far, HapSolo2 converted duplicated orthologs to single-copy orthologs at a cost of at most 1.3 percentage points of completeness.

- *A. funestus*. D fell from 7.8% to 2.8% and S rose from 88.1% to 92.9% (RW, ODB12.2), with C at 95.7% against 95.9% unreduced. All three releases gave the same C (95.7%) and scores within 1.5 points of one another (181.5 to 183.0).
- *P. americana*. D fell from 52.0% to 5.6% and S nearly doubled, from 47.6% to 92.6% (SA, ODB10), with C at 98.3% against 99.6%. The three releases differ by at most 0.2 points of the score; ODB10 and ODB12.2 tie at 179.7, and ODB10 is selected for its higher C.
- *V. vinifera*. D fell from 17.9% to 10.4% and S rose from 80.7% to 87.8% (SA, ODB12.2), with C at 98.1% against 98.6%. All three releases are within 0.1 points (165.0 to 165.1), although the ODB9 run retains 112 contigs and the ODB10 and ODB12.2 runs 47 to 53.
- *A. radiata*. Two of six candidates have been assessed. The actinopterygii ODB10 SA run reduced D from 23.2% to 20.7% (score 121.1 against 114.5 unreduced). The actinopterygii ODB9 run on the complete cache (PID 0.6112, 11,572 contigs) reproduces exactly the contig set of an earlier run of the same thresholds, which BUSCO assessed at C 91.7%, S 81.5% and D 10.2% (score 152.8).

**Table 3:** Independent BUSCO assessment (v6.1, genome mode, odb12.2) of the best run per training release. Score = *C* + *S-*2*D* in BUSCO percentages. One validation lineage per species (Table 1), independent of the training release. Bold marks the highest score among the logged runs assessed so far. ‡ Earlier run not in the timed logs (optimizer and code version unrecorded); the *A. radiata* run at PID 0.6112 cannot be reproduced from the alignment cache on disk.

| Run | Training lineage | Contigs | C (%) | S (%) | D (%) | F (%) | M (%) | Score |
| --- | --- | --- | --- | --- | --- | --- | --- | --- |
| Unreduced | n/a | 1,073 | 95.9 | 88.1 | 7.8 | 0.2 | 3.9 | 168.4 |
| SD, ODB9 | diptera | 903 | 95.7 | 92.7 | 2.9 | 0.2 | 4.1 | 182.6 |
| SA, ODB10 | diptera | 879 | 95.7 | 92.4 | 3.3 | 0.2 | 4.1 | 181.5 |
| **RW, ODB12.2** | diptera | 854 | **95.7** | **92.9** | **2.8** | 0.2 | 4.1 | **183.0** |
| Earlier run‡, ODB12.2 | diptera | 908 | 95.7 | 92.5 | 3.2 | 0.2 | 4.1 | 181.8 |

**(b) *Persea americana* cv. Gwen**
| Run | Training lineage | Contigs | C (%) | S (%) | D (%) | F (%) | M (%) | Score |
| --- | --- | --- | --- | --- | --- | --- | --- | --- |
| Unreduced | n/a | 5,122 | 99.6 | 47.6 | 52.0 | 0.0 | 0.4 | 43.2 |
| RW, ODB9 | embryophyta | 496 | 98.3 | 92.6 | 5.7 | 0.1 | 1.6 | 179.5 |
| **SA, ODB10** | embryophyta | 487 | **98.3** | **92.6** | **5.6** | 0.1 | 1.6 | **179.7** |
| SA, ODB12.2 | embryophyta | 477 | 98.1 | 92.6 | 5.5 | 0.1 | 1.8 | 179.7 |
| Earlier run‡, ODB12.2 | embryophyta | 687 | 98.1 | 92.5 | 5.6 | 0.1 | 1.7 | 179.4 |

**(c) *Vitis vinifera* cv. Chardonnay**
| Run | Training lineage | Contigs | C (%) | S (%) | D (%) | F (%) | M (%) | Score |
| --- | --- | --- | --- | --- | --- | --- | --- | --- |
| Unreduced | n/a | 684 | 98.6 | 80.7 | 17.9 | 0.2 | 1.3 | 143.5 |
| RW, ODB9 | embryophyta | 112 | 98.1 | 87.7 | 10.4 | 0.2 | 1.6 | 165.0 |
| SA, ODB10 | embryophyta | 53 | 98.1 | 87.7 | 10.4 | 0.2 | 1.6 | 165.0 |
| **SA, ODB12.2** | embryophyta | 47 | **98.1** | **87.8** | **10.4** | 0.2 | 1.6 | **165.1** |
| Earlier run‡, ODB12.2 | embryophyta | 67 | 98.1 | 87.8 | 10.4 | 0.2 | 1.6 | 165.1 |

**(d) *Amblyraja radiata***
| Run | Training lineage | Contigs | C (%) | S (%) | D (%) | F (%) | M (%) | Score |
| --- | --- | --- | --- | --- | --- | --- | --- | --- |
| Unreduced | n/a | 16,218 | 92.0 | 68.9 | 23.2 | 2.8 | 5.2 | 114.5 |
| SD, ODB9 | actinopterygii | 16,135 | pending |  |  |  |  |  |
| **SA, ODB10** | actinopterygii | 16,037 | **91.6** | **70.9** | **20.7** | 2.8 | 5.5 | **121.1** |
| SA, ODB12.2 | actinopterygii | 15,884 | pending |  |  |  |  |  |
| SA, ODB9 | metazoa | 15,771 | pending |  |  |  |  |  |
| SD, ODB10 | metazoa | 15,270 | pending |  |  |  |  |  |
| RW, ODB12.2 | metazoa | 15,138 | pending |  |  |  |  |  |
| Earlier run‡, ODB9 | actinopterygii | 11,572 | 91.7 | 81.5 | 10.2 | 2.9 | 5.4 | 152.8 |

Two observations bear on the choice of training release. First, the internal ranking did not predict the BUSCO ranking: for *A. funestus*, the ODB12.2 run ranked last of three on the internal score and first on BUSCO. Second, when the training release is ODB12.2 and the validation lineage is from the same release, training and validation share a gene set; in the three species assessed, the ODB12.2 runs lead the best ODB9 or ODB10 run by at most 0.4 points, so this overlap does not change any conclusion here.

### Agreement among optimizers and OrthoDB releases

Table 4 compares the sets of contigs retained by RW and SD across training releases. For *A. funestus*, *P. americana* and *V. vinifera* the sets were reconstructed from each run’s thresholds (for the 12 *A. funestus* and *P. americana* runs, the reconstruction reproduces 11 outputs exactly and the remaining one within one contig); for *A. radiata* they were read from the primary assemblies of the runs on the complete alignment cache, which the reconstruction reproduces exactly for 8 of 12 runs and within two contigs for the remaining 4. RW and SD retain nearly identical sets: the contigs kept by all three releases under RW and under SD overlap with Jaccard indices of 0.923 to 0.991. Across releases, agreement is high for *A. funestus* (0.932 to 0.944) and *P. americana* (0.834 to 0.886), intermediate for *A. radiata* with actinopterygii (0.771 to 0.806), and low for *V. vinifera* (0.425 to 0.464) and for *A. radiata* with metazoa (0.229 to 0.232). For *V. vinifera*, the ODB9 runs keep about twice as many contigs as the ODB10 and ODB12.2 runs without a difference in the BUSCO assessment (Table 3). For *A. radiata* with metazoa, the ODB9 runs keep 11,564 to 11,739 contigs against 2,697 to 3,210 for ODB10 and ODB12.2, and 8,441 (RW) and 8,720 (SD) contigs are kept by ODB9 alone (Figure 4).

**Figure 4:**
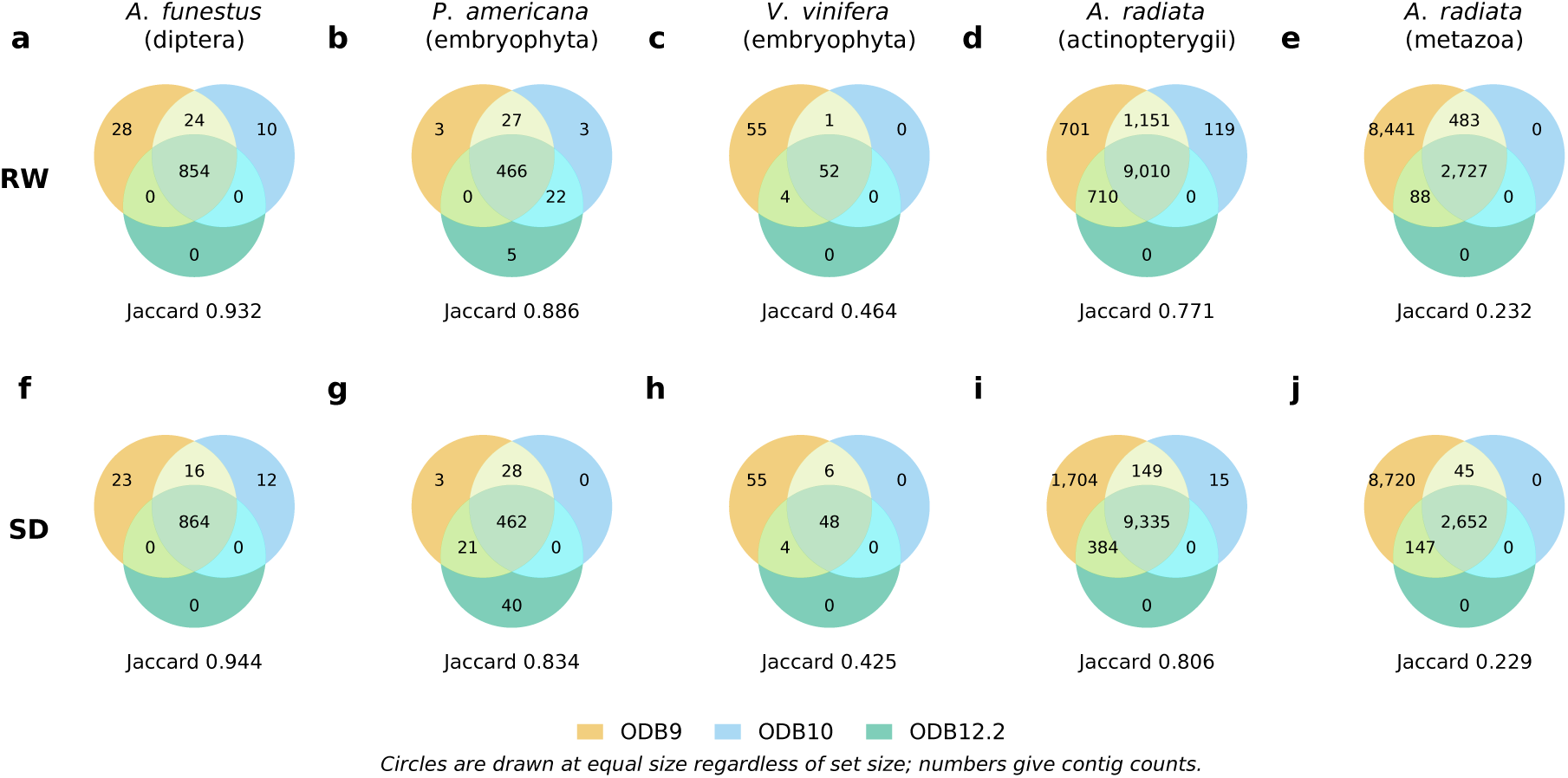
Overlap of the contigs kept by the ODB9, ODB10 and ODB12.2 runs for each species and training lineage, with RW (a to e) and SD (f to j) (Table 4). Numbers give contig counts in each region; circles are drawn at equal size regardless of set size, so that panels of 52 and 11,739 contigs read alike. The Jaccard index of the three sets (contigs kept by all three releases divided by contigs kept by any) is given under each diagram. *A. radiata* sets are from the runs on the complete alignment cache.

**Table 4:** Agreement of kept-contig sets across training releases and between optimizers. Kept sets reconstructed from each run’s thresholds, except *A. radiata*, whose sets are read from the primary assemblies of the runs on the complete alignment cache. Core = contigs kept by all three releases for that optimizer.

**(a) Across releases, per optimizer**
| Species (lineage) | Optimizer | Kept, ODB9 | Kept, ODB10 | Kept, ODB12.2 | Core | Union | Jaccard |
| --- | --- | --- | --- | --- | --- | --- | --- |
| *A. funestus* (diptera) | RW | 906 | 888 | 854 | 854 | 916 | 0.932 |
| *A. funestus* (diptera) | SD | 903 | 892 | 864 | 864 | 915 | 0.944 |
| *P. americana* (embryophyta) | RW | 496 | 518 | 493 | 466 | 526 | 0.886 |
| *P. americana* (embryophyta) | SD | 514 | 490 | 523 | 462 | 554 | 0.834 |
| *V. vinifera* (embryophyta) | RW | 112 | 53 | 56 | 52 | 112 | 0.464 |
| *V. vinifera* (embryophyta) | SD | 113 | 54 | 52 | 48 | 113 | 0.425 |
| *A. radiata* (actinopterygii) | RW | 11,572 | 10,280 | 9,720 | 9,010 | 11,691 | 0.771 |
| *A. radiata* (actinopterygii) | SD | 11,572 | 9,499 | 9,719 | 9,335 | 11,587 | 0.806 |
| *A. radiata* (metazoa) | RW | 11,739 | 3,210 | 2,815 | 2,727 | 11,739 | 0.232 |
| *A. radiata* (metazoa) | SD | 11,564 | 2,697 | 2,799 | 2,652 | 11,564 | 0.229 |

**(b) RW core against SD core**
| Species (lineage) | RW core | SD core | Shared | Jaccard |
| --- | --- | --- | --- | --- |
| *A. funestus* (diptera) | 854 | 864 | 854 | 0.988 |
| *P. americana* (embryophyta) | 466 | 462 | 462 | 0.991 |
| *V. vinifera* (embryophyta) | 52 | 48 | 48 | 0.923 |
| *A. radiata* (actinopterygii) | 9,010 | 9,335 | 8,891 | 0.940 |
| *A. radiata* (metazoa) | 2,727 | 2,652 | 2,650 | 0.971 |

### Comparison with the original HapSolo

On *P. americana* with ODB9 (Table 5), the original HapSolo and a CPU-only refactor of HapSolo2 reached the same cost (0.2200), retaining 1,083 and 903 contigs with 15.9% and 15.8% internal duplication. The GPU runs of 100,000 iterations reached a cost of 0.1036 to 0.1045, retaining 496 to 618 contigs with 5.4% to 5.5% internal duplication and a contig N50 of 4,601 to 4,608 kb against 3,366 kb. The original HapSolo run used 50,000 iterations (Solares et al. 2023) and BUSCO v3 ortholog calls, against 100,000 iterations over 84 walkers and the miniprot ortholog table for HapSolo2; both used a lower bound of 0.2 on each threshold, so the difference reflects the iteration budget and the ortholog table rather than the search space.

**Table 5:** Original HapSolo and HapSolo2 on *P. americana* cv. Gwen (ODB9, internal scores). Legacy rows were produced by earlier code and are kept for comparison.

| Run | PID | QRPct | QPct | Contigs | Size (Mb) | N50 (kb) | C (%) | S (%) | D (%) | F (%) | M (%) | Cost |
| --- | --- | --- | --- | --- | --- | --- | --- | --- | --- | --- | --- | --- |
| Original HapSolo (legacy) | 0.6206 | 0.9824 | 0.2010 | 1,083 | 1,041.7 | 3,366 | 97.0 | 81.1 | 15.9 | 1.0 | 1.9 | 0.2200 |
| HapSolo2 CPU refactor (legacy) | 0.3418 | 0.6706 | 0.2023 | 903 | 1,027.4 | 3,387 | 96.9 | 81.1 | 15.8 | 1.0 | 2.1 | 0.2200 |
| HapSolo2 GPU RW, 100k | 0.2214 | 0.2792 | 0.3660 | 496 | 860.5 | 4,608 | 95.2 | 89.8 | 5.4 | 0.9 | 3.9 | 0.1036 |
| HapSolo2 GPU SD, 100k | 0.2557 | 0.2246 | 0.3663 | 514 | 862.5 | 4,608 | 95.2 | 89.8 | 5.4 | 0.9 | 3.9 | 0.1036 |
| HapSolo2 GPU SA, 100k | 0.4355 | 0.5177 | 0.3657 | 618 | 871.7 | 4,601 | 95.2 | 89.7 | 5.5 | 0.9 | 3.9 | 0.1045 |

### Computational performance

#### GPU against CPU for RW and SD

On *P. americana* (5,122 contigs, 1,440 orthologs, 49,285 alignment records), timing the optimization phase only with five replicates per configuration, 84 GPU walkers on one RTX A6000 were compared with 10 CPU processes (Table 6; Figure 5). The GPU speedup grew with the iteration budget as fixed launch costs were amortized, from 1.1-fold at 1,000 RW iterations to 5.6-fold at 100,000, and was larger for SD (3.6- to 7.6-fold), which submits eight candidates per walker per iteration. GPU timings were also far more consistent: at 100,000 SD iterations the coefficient of variation was 0.3% on the GPU and 18.9% on the CPU, where the slowest pool worker gates each batch. This benchmark was run on 2026-07-01 on an earlier ortholog table, and only its timings are reported here.

**Figure 5:**
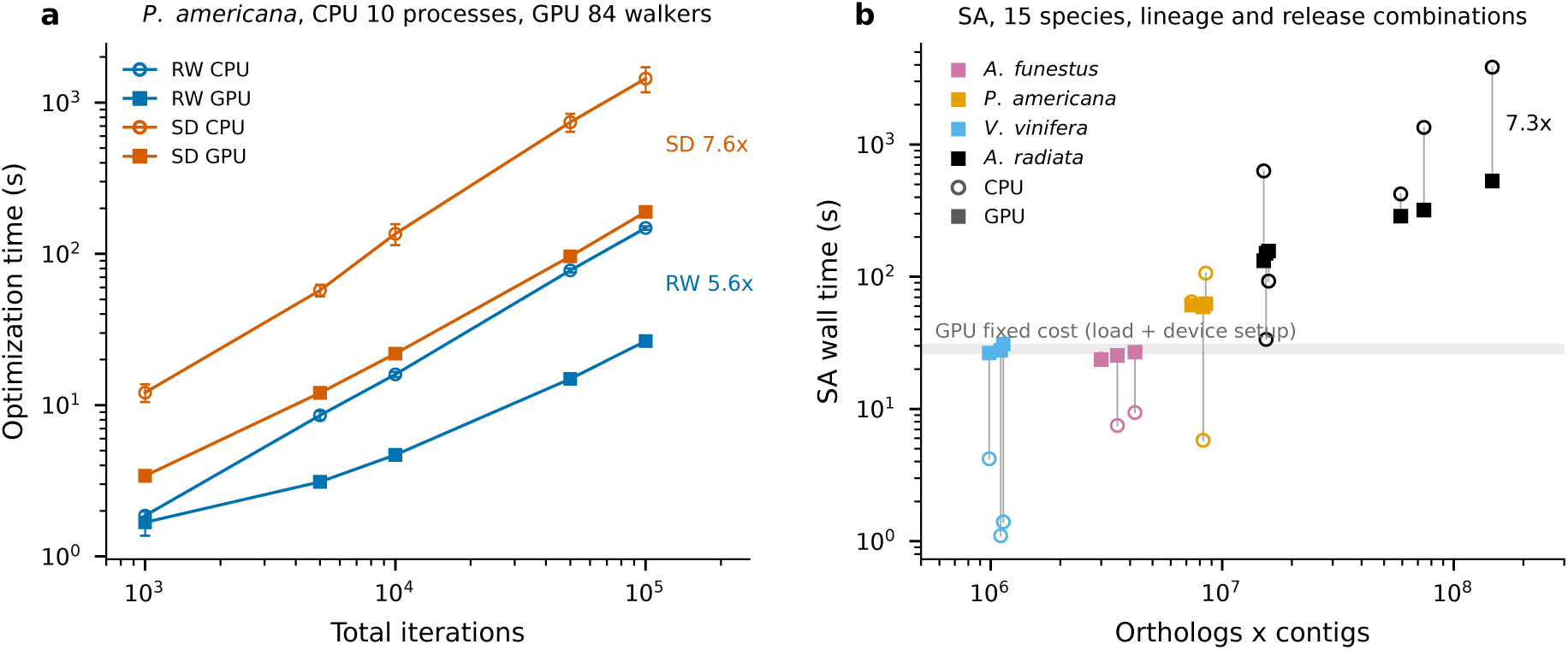
Computational performance. (a) Optimization time of RW and SD on *P. americana* by total iteration budget on the CPU (10 processes, open circles) and the GPU (84 walkers on one RTX A6000, filled squares); mean and standard deviation of five replicates (Table 6). Labels give the CPU to GPU speedup at 100,000 iterations. (b) SA wall time, end to end including data loading, on the CPU and the GPU against the product of orthologs and contigs for the 15 species, lineage and release combinations (Table 8). CPU points (open circles) are the fastest configuration that reached the best cost among those tested; GPU points (filled squares) are the 100,000-iteration runs. Gray segments join the two points of one combination, and the gray band marks the GPU fixed cost of loading data and setting up the device (26 to 31 s). The label gives the speedup for the largest problem, *A. radiata* with actinopterygii ODB12.2. *A. radiata* timings in (b) were measured on an alignment cache of 1,502 of the 207,119 alignments.

**Table 6:**
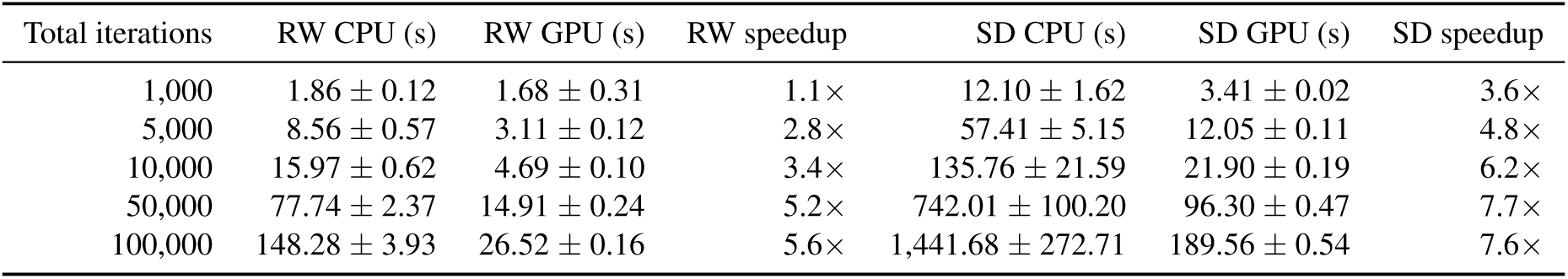
Optimization time of RW and SD on CPU and GPU for *P. americana* (mean ± s.d., *n* = 5). CPU: 10 processes. GPU: 84 walkers on one RTX A6000. Data loading and FASTA output excluded.

#### SA convergence and CPU against GPU

Table 7 reports the best cost reached by GPU SA at budgets of 1,000 to 100,000 total iterations, and Table 8 compares it with CPU SA at 1 or 4 processes and 1,000 to 10,000 iterations per process. On the GPU, 9 of the 15 species, lineage and release combinations reached their 100,000-iteration cost by 10,000 iterations, and the costs are not monotone in the budget (for *P. americana* ODB9, 0.103635 at 10,000 and 50,000 iterations against 0.104489 at 100,000), reflecting run-to-run variation between unseeded single runs. The CPU and the GPU reached costs within 2.0% of each other in all 15 combinations and identical costs in 9. The GPU was faster for the largest problems (*A. radiata*, actinopterygii ODB12.2, 9,048 orthologs by 16,218 contigs: 529.7 s against 3,845.9 s, 7.3-fold) but slower for the smallest, where the fixed cost of loading data and setting up the device (wall time 26.5 to 31.0 s for *V. vinifera*, of which 8.9 to 11.0 s is optimization) exceeded the entire CPU run (*V. vinifera* ODB10: 27.7 s against 1.1 s). The crossover lies near 5,000 contigs: *P. americana* (5,122 contigs) ran 1.8-to 2.1-fold faster on the GPU for the two releases whose CPU runs needed 10,000 iterations per process, and slower for the one that converged at 1,000.

**Table 7:** GPU SA convergence: best cost by total iteration budget (84 walkers, RTX A6000). Bold marks the first budget that reaches the 100,000-iteration cost. Opt = optimization time only; Wall = end to end, including data loading.

| Species, lineage, release | 1k | 5k | 10k | 50k | 100k | Opt, 100k (s) | Wall, 100k (s) |
| --- | --- | --- | --- | --- | --- | --- | --- |
| *A. funestus*, diptera, ODB9 | 0.053558 | 0.053175 | 0.052752 | **0.052350** | 0.052350 | 16.3 | 23.6 |
| *A. funestus*, diptera, ODB10 | 0.109422 | 0.109048 | **0.108674** | 0.108674 | 0.108674 | 18.2 | 25.4 |
| *A. funestus*, diptera, ODB12.2 | 0.099913 | **0.099592** | 0.099884 | 0.099592 | 0.099592 | 19.6 | 26.9 |
| *P. americana*, embryophyta, ODB9 | **0.104489** | 0.104489 | 0.103635 | 0.103635 | 0.104489 | 26.4 | 61.0 |
| *P. americana*, embryophyta, ODB10 | 0.336093 | **0.334988** | 0.334988 | 0.334988 | 0.334988 | 27.2 | 59.1 |
| *P. americana*, embryophyta, ODB12.2 | 0.440945 | 0.438811 | 0.439685 | **0.437173** | 0.437173 | 28.7 | 62.5 |
| *V. vinifera*, embryophyta, ODB9 | **0.039330** | 0.039330 | 0.039330 | 0.038574 | 0.039330 | 9.0 | 26.5 |
| *V. vinifera*, embryophyta, ODB10 | **0.222727** | 0.222727 | 0.222727 | 0.222727 | 0.222727 | 8.9 | 27.7 |
| *V. vinifera*, embryophyta, ODB12.2 | **0.265432** | 0.265432 | 0.265432 | 0.265432 | 0.265432 | 11.0 | 31.0 |
| *A. radiata*, actinopterygii, ODB9 | 0.303706 | 0.303706 | 0.303706 | 0.303222 | **0.302921** | 217.3 | 319.7 |
| *A. radiata*, actinopterygii, ODB10 | 0.456000 | 0.455418 | **0.454836** | 0.454836 | 0.454836 | 187.8 | 287.2 |
| *A. radiata*, actinopterygii, ODB12.2 | 0.458552 | 0.458173 | 0.458173 | 0.457878 | **0.457287** | 425.1 | 529.7 |
| *A. radiata*, metazoa, ODB9 | 0.317529 | 0.315638 | 0.315638 | 0.315638 | **0.313754** | 62.9 | 156.6 |
| *A. radiata*, metazoa, ODB10 | 3.042373 | **2.975000** | 3.008403 | 2.975000 | 2.975000 | 63.2 | 150.6 |
| *A. radiata*, metazoa, ODB12.2 | 1.035806 | 1.017903 | 1.023018 | 1.015306 | **1.015228** | 57.1 | 132.1 |

**Table 8:** SA on CPU and GPU: best cost and wall time. CPU: best over the tested configurations (1 or 4 processes × 1,000, 5,000 or 10,000 iterations per process), with the wall time of the fastest configuration reaching that cost. GPU: 100,000 total iterations.

| Species, lineage, release | CPU best | CPU configuration | CPU wall (s) | GPU best | GPU wall (s) | GPU/CPU cost |
| --- | --- | --- | --- | --- | --- | --- |
| *A. funestus*, diptera, ODB9 | 0.052752 | 1 × 5k | 23.8 | 0.052350 | 23.6 | 0.9924 |
| *A. funestus*, diptera, ODB10 | 0.108674 | 4 × 1k | 7.5 | 0.108674 | 25.4 | 1.0000 |
| *A. funestus*, diptera, ODB12.2 | 0.099592 | 4 × 1k | 9.4 | 0.099592 | 26.9 | 1.0000 |
| *P. americana*, embryophyta, ODB9 | 0.103635 | 4 × 10k | 64.6 | 0.104489 | 61.0 | 1.0082 |
| *P. americana*, embryophyta, ODB10 | 0.334988 | 1 × 1k | 5.8 | 0.334988 | 59.1 | 1.0000 |
| *P. americana*, embryophyta, ODB12.2 | 0.437173 | 4 × 10k | 106.5 | 0.437173 | 62.5 | 1.0000 |
| *V. vinifera*, embryophyta, ODB9 | 0.038574 | 1 × 5k | 4.2 | 0.039330 | 26.5 | 1.0196 |
| *V. vinifera*, embryophyta, ODB10 | 0.222727 | 1 × 1k | 1.1 | 0.222727 | 27.7 | 1.0000 |
| *V. vinifera*, embryophyta, ODB12.2 | 0.265432 | 4 × 1k | 1.4 | 0.265432 | 31.0 | 1.0000 |
| *A. radiata*, actinopterygii, ODB9 | 0.302529 | 4 × 10k | 1,347.6 | 0.302921 | 319.7 | 1.0013 |
| *A. radiata*, actinopterygii, ODB10 | 0.454836 | 4 × 5k | 422.7 | 0.454836 | 287.2 | 1.0000 |
| *A. radiata*, actinopterygii, ODB12.2 | 0.457895 | 4 × 5k | 3,845.9 | 0.457287 | 529.7 | 0.9987 |
| *A. radiata*, metazoa, ODB9 | 0.313754 | 4 × 10k | 92.7 | 0.313754 | 156.6 | 1.0000 |
| *A. radiata*, metazoa, ODB10 | 2.975000 | 4 × 1k | 33.6 | 2.975000 | 150.6 | 1.0000 |
| *A. radiata*, metazoa, ODB12.2 | 1.017632 | 4 × 10k | 631.5 | 1.015228 | 132.1 | 0.9976 |

#### Optimizer cost

Under the same GPU protocol, SD needed 2.7 to 6.5 times the wall time of RW in the 12 logged run pairs (Table 2), due to its up to eight evaluations per iteration, and it reached a strictly better optimum than RW and SA in only 3 of 15 combinations, by at most 1.7%. SA wall times were within 16% of RW in the same 12 combinations.

## Discussion

We have presented HapSolo2, a reimplementation of HapSolo that keeps the original formulation of haplotig removal as threshold optimization and replaces every part of the implementation that limited its use: the per-contig BUSCO runs, the Python 2.7 and pandas evaluation loop, the single optimizer, and the separate post-processing. The resulting pipeline runs from a draft assembly to primary and secondary assemblies with one command, in under a minute of optimization for assemblies of a few thousand contigs and in under ten minutes for a 3.2 Gb assembly of 16,218 contigs scored against 9,048 orthologs. On the four assemblies tested, the reduced assemblies retained 91.6% to 98.3% complete orthologs under an independent BUSCO assessment while duplicated orthologs fell by 2.5 to 46.4 percentage points, and the largest gain came from the most duplicated assembly, the avocado, whose single-copy fraction nearly doubled.

The clearest result of the optimizer comparison is that it does not matter much which optimizer is used. Random forward walk, steepest descent and simulated annealing reached best costs within 1.8% of one another in all 15 species, lineage and release combinations, and the sets of contigs they retained overlapped with Jaccard indices above 0.92. The threshold triples at those optima, however, differed widely, with PID ranging from 0.22 to 0.44 among three runs that retained nearly the same contigs. We conclude that the cost landscape consists of broad basins in which many threshold triples remove the same ortholog-bearing contigs, so that the search is far less sensitive to the optimizer than we assumed when we proposed simulated annealing as an area for growth (Solares et al. 2021). A practical consequence is that the thresholds reported by a run should be read as one representative of a basin rather than as a precise setting, and that the kept set, not the triple, is the result to compare between runs. A second consequence is a recommendation: random forward walk on the GPU is the default, since it reached the same optimum as steepest descent at a quarter to a sixth of the wall time, and steepest descent is worth its cost only where a strictly lower optimum matters, as it did by at most 1.7% on *A. radiata*.

The choice of ortholog dataset matters more than the choice of optimizer, but not in the way the internal cost suggests. Internal costs are not comparable across OrthoDB releases, since each release defines a different gene set and ODB10 classifies no ortholog as fragmented, and the internal ranking of releases did not predict the ranking under an independent BUSCO assessment. After that assessment, however, the three releases gave the same completeness and scores within 1.5 points of one another in every species assessed, and the contigs on which they disagreed carried few or no orthologs. For new analyses we recommend ODB12.2, which is the curated current release, and we advise against ODB12 (OrthoDB 12.1), which lacks length cutoffs and produced a near-meaningless classification. For reproducing results obtained with the original HapSolo, the archived ODB9 datasets remain usable. In either case the reduced assembly should be assessed with a BUSCO lineage that was not used for training, with the caveat that the ODB12.2 validation lineages share a gene set with ODB12.2 training; in our data this overlap was worth at most 0.4 points of the score.

Taxa without a dedicated lineage require more care. For the thorny skate, a cartilaginous fish, the actinopterygii datasets removed 1.3% to 6.3% of the assembly and the metazoa datasets 8.1% to 24.3%, and the metazoa ODB10 and ODB12.2 runs left 40.6% to 69.9% of their few hundred orthologs duplicated while removing a fifth of the assembly. A small, distant gene set gives the cost little purchase on the contigs that matter, and the optimizer then removes sequence that the cost is indifferent to. BUSCO assessment of the six *A. radiata* candidates on the complete alignment cache is in progress; until it is complete we recommend the larger and closer lineage (actinopterygii) and independent validation of the reduced assembly.

The GPU implementation changes what is practical rather than what is possible. The CPU and GPU paths evaluate identical candidates and reached costs within 2.0% of each other in all 15 SA comparisons, so the GPU adds speed only. That speed depends on problem size: the fixed cost of loading data and initializing the device, 27 to 31 s for the smallest assembly, exceeds an entire CPU run for assemblies of a few hundred contigs, while the largest problem ran 7.3-fold faster on the GPU, and the crossover lies near 5,000 contigs. The speedup also depends on how much work each iteration offers the device, which is why steepest descent, with up to eight candidates per walker per iteration, gained more from the GPU (7.6-fold) than random walk (5.6-fold) at the same budget. The practical value of the GPU is therefore for gigabase-scale assemblies with tens of thousands of contigs, where the original HapSolo needed 13 h 45 min for the thorny skate at 50,000 iterations on 10 cores (Solares et al. 2021) and HapSolo2 needs 1 to 7 min of optimization (2 to 9 min end to end) on one workstation GPU, and for the many-walker, large-budget runs that make the choice of optimizer and starting point immaterial. On the same avocado assembly and ODB9 gene set, HapSolo2 reached a cost of 0.1036 against 0.2200 for the original HapSolo and retained 496 rather than 1,083 contigs, with internal duplication falling from 15.9% to 5.4% and N50 rising from 3,366 to 4,608 kb. Both implementations bound each threshold below at 0.2, so the improvement reflects the larger iteration budget, distributed over 84 walkers, and the ortholog table.

Three limitations follow from the results. First, the cost is defined only on contigs that carry orthologs, so contigs without a conserved gene are invisible to it, and the optimizer keeps or removes them according to the alignment thresholds alone. The low agreement among releases on *V. vinifera*, where the ODB9 runs kept about twice as many contigs as the others at the same BUSCO score, shows that these contigs can be numerous in a well-assembled genome. Second, HapSolo2 still uses no read-coverage information, the point of departure from PurgeHaplotigs and purge_dups that we identified previously, and the thresholds are global, so a region whose haplotypes are more divergent than the rest of the genome is purged more conservatively than its neighbors. Third, the benefit is bounded by the duplication present: for *A. funestus*, with 7.8% duplicated orthologs, the gain was 5.0 percentage points, and we expect none for genomes near the 0.33% heterozygosity and 2.1% duplicated BUSCOs of *Peromyscus leucopus*, for which we previously found no benefit and suggested roughly 5% duplicated BUSCOs as a lower limit for the method (Solares et al. 2021). Two further points concern reproducibility. The GPU random-walk and simulated-annealing runs are not seeded by default, and the SA sweep shows run-to-run variation of up to 2.0% in cost between single runs, so seeded replicates are advisable when thresholds are reported. The results for *A. funestus*, *P. americana* and *V. vinifera* were produced with the July 2026 code, whose SA reheat differs from the released code in the handling of plateaus; the *A. radiata* results were produced with the released code.

That is not to say that HapSolo2 cannot be improved further. We can see three areas for growth. The first remains read coverage, which could be added as a fourth filter on the same batched evaluation at little cost once per-contig depth is tabulated. The second is a per-contig term in the cost for contigs without orthologs, for example an over-representation of shared *k*-mers, so that the optimizer is no longer indifferent to them. The third is the scaffolding step itself: the contig order and the ortholog table that HapSolo2 already holds could inform the removal of haplotigs that Hi-C scaffolding would otherwise join, and scaffolding the reduced assemblies, which we showed previously to benefit from haplotig removal (Solares et al. 2021), will show how much of that benefit the current reduction already captures.

## Conclusions

HapSolo2 retains the approach of HapSolo, haplotig removal by optimization of alignment thresholds against an ortholog-based cost, and makes it practical: one command runs the pipeline from a draft assembly to primary and secondary assemblies, ortholog classification takes one miniprot run instead of one BUSCO run per contig, and 100,000 optimizer iterations complete in seconds to minutes on one GPU, 5.6-to 7.6-fold faster than on 10 CPU processes. Across four diploid assemblies spanning 212 Mb to 3.2 Gb and three OrthoDB releases, the three optimizers reached the same optima to within 1.8%, which makes the fast random forward walk the default and the retained contig set, rather than the threshold triple, the result to report. Under an independent BUSCO assessment the reduced assemblies lost at most 1.3 percentage points of completeness while duplicated orthologs fell from 52.0% to 5.6% in the most heterozygous assembly. Internal costs are not comparable across OrthoDB releases, so we recommend training on ODB12.2 and validating with a lineage not used for training; for taxa without a dedicated lineage, the larger and closer dataset should be preferred and the result validated before use. HapSolo2 is available as a pip-installable package, as an Apptainer image with pinned tools and the CUDA runtime, and with archived copies of the ODB9 datasets, so that both new analyses and the results of the original HapSolo remain reproducible.

## Methods

### Species and data

The *A. funestus* contig assembly (NCBI BioProject PRJNA494870; Ghurye et al. 2019) and the *A. radiata* contig assembly (GenBank GCA_010909765.1, the sAmbRad1 primary haplotype assembled by the Vertebrate Genomes Project; Rhie et al. 2021) are the files analyzed in Solares et al. (2021). Both are FALCON-Unzip primary contig assemblies from PacBio data (240× and 128× coverage, respectively) (Solares et al. 2021). The *P. americana* cv. Gwen contig assembly (5,122 contigs, 1,449.9 Mb) is the assembly of Solares et al. (2023) before haplotig removal: PacBio Sequel reads were assembled with Canu v2.1 and polished with two passes of GenomicConsensus v2.33 and two passes of Pilon v1.23 with 19× Illumina coverage. The *V. vinifera* cv. Chardonnay contig assembly (684 contigs, 606.0 Mb, contig N50 24.5 Mb) is the Chardonnay genome assembly of Zhou et al. (2019), NCBI BioProject PRJNA550461, also the *V. vinifera* data source of Solares et al. (2021). *Culex tarsalis* data were analyzed separately and are not reported here.

### Preprocessing

Assemblies were preprocessed with `hapsolo-cli preprocess`. The first whitespace-delimited token of each FASTA header was kept, every character outside `[a-zA-Z0-9.]` was replaced by an underscore, and headers that became identical were truncated to the shortest prefix length at which all names remained unique. The sanitized assembly was written with one sequence line per contig, individual contig files were written for contigs shorter than 10 Mb, and the mapping from original to sanitized names was recorded.

### Self-alignment

Each sanitized assembly was aligned to itself with minimap2 2.31 (Li 2018) with the parameters set by

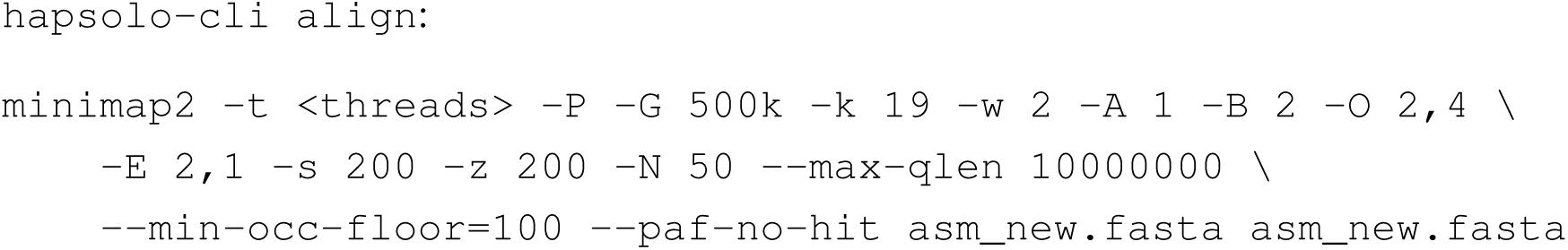

Option `-P` retains all chains rather than selecting primary chains, so that every overlap between contigs is reported; with `-P` in effect, `-N` has no effect. The small k-mer and window sizes (`-k 19 -w 2`) increase sensitivity to divergent haplotigs, and the low mismatch and gap penalties tolerate the differences expected between haplotypes. The `-x asm5` preset, which is tuned for high-identity assembly-to-reference alignment, discards most haplotig candidates and should not be used. The PAF output was compressed with gzip and parsed once into the `.hap` cache with `hapsolo-cli cache` (minimum query contig length 1,000 bp; PID, QPct and QRPct at least 0.2). BLAT (`blat asm asm out.psl-noHead`) is supported as an alternative aligner but was not used for the reported results.

### Ortholog datasets and classification

Ortholog groups were taken from the BUSCO lineage datasets of OrthoDB releases 9, 10 and 12.2 (Kriventseva et al. 2019; Manni et al. 2021; Tegenfeldt et al. 2025): diptera for *A. funestus* (2,799, 3,285 and 3,914 groups, respectively), embryophyta for *P. americana* and *V. vinifera* (1,440, 1,614 and 1,658 groups), and both actinopterygii (4,584, 3,640 and 9,048 groups) and metazoa (978, 954 and 932 groups) for *A. radiata*. ODB10 and ODB12.2 datasets were downloaded from the BUSCO data archive (https://busco-data.s3.amazonaws.com/lineages/) diptera_odb9, metazoa_odb9 2016-10-21; embryophyta_odb9, actinopterygii_odb9 2016-02-13; all ODB10 datasets 2024-01-08; all ODB12.2 datasets 2026-05-13 and ODB9 datasets from our Zenodo archive (Zdobnov et al. 2026, https://doi.org/10.5281/zenodo.20987183). ODB12 (OrthoDB 12.1) was evaluated on *P. americana* and rejected, since its datasets lack `lengths_cutoff` and their group identifiers are not retained in ODB12.2 (BUSCO development team 2026).

Ortholog proteins (the `ancestral_variants` set for ODB9 and `refseq_db.faa` for ODB10 and ODB12.2) were aligned to each assembly with miniprot 0.18 (Li 2023) in PAF mode with `-I` (maximum intron length 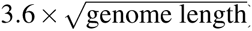). For ODB10 and later, protein identifiers were mapped to ortholog groups by trying underscore-delimited prefixes of the identifier, shortest first, against the keys of `scores_cutoff`, and proteins whose group was absent from `scores_cutoff` were excluded. For each group and contig the highest-scoring alignment was retained; an alignment was significant if its miniprot score (`AS:i`) met the group’s `scores_cutoff` value, Complete if its aligned protein length reached the group’s expected length from `lengths_cutoff` (two scale units below the group’s central length, as above) and Fragmented otherwise, and a group with no significant alignment on any contig was Missing. The ortholog tables used for the reported results were generated on 2026-07-09 and 2026-07-10 with the July 2026 classifier, which applied only the score test above, used a fallback expected length of 0.95 times the longest hit where `lengths_cutoff` was absent, and ran miniprot with `-B 5 -j 1`. The released classifier additionally requires a score-to-cutoff ratio of at least 2.5 and an alignment coverage of at least 0.7, counts a group on a second contig only when that contig’s score is at least 0.9 of the best, uses a 0.85 fallback, and runs miniprot with `-B 25 -j 2`, so ortholog tables regenerated with the released code differ from those used here.

### Optimization runs

For each species, training lineage and release, HapSolo2 was run on the GPU with RW, SD and SA for 100,000 total iterations (84 walkers × 1,191 iterations per walker, 100,044 evaluations) with the default cost weights (θ*_S_* = θ*_D_*= θ*_M_*= 1, θ*_F_*= 0), default search space ([0.2, 1] in each of PID, QPct and QRPct), step size and plateau resolution of 1 × 10^−4^, plateau window of 10 iterations, and *B* = 1, giving 45 runs (four species, five species-lineage combinations, three releases, three optimizers). SA used the data-calibrated initial temperature, the geometric schedule with α set from 1,191 iterations per walker, and *T*_min_ = 1 ×10^−6^. RW and SD runs were timed end to end, including loading of the alignment cache and ortholog table; the *V. vinifera* RW and SD runs were not in the timed logs and their wall times are taken from the results table (marked † in Table 2). SA convergence was measured at 1,000, 5,000, 10,000, 50,000 and 100,000 total iterations on the GPU, and on the CPU with 1 or 4 processes at 1,000, 5,000 and 10,000 iterations per process; each configuration was a single unseeded run. All reported runs used the batched GPU evaluation path with host-side walker logic July 2026 code, preserved as `archive/hapsolo_pip_install_2026-07-21/` in the repository.

### Independent validation

Reduced assemblies were assessed with BUSCO v6.1 (Tegenfeldt et al. 2025) in genome mode, with its default miniprot-based gene mapping, against a single odb12.2 lineage per species: diptera for *A. funestus*, embryophyta for *P. americana* and *V. vinifera*, and vertebrata for *A. radiata*, for which neither training lineage is a close match. The assessment therefore shares the protein aligner with HapSolo2’s classifier but not the lineage dataset or the classification rules. For each species, training lineage and release, the run with the highest *C* + *S* − 2*D* on the internal scores was assessed (ties broken by higher C, then higher S, then fewer contigs), and the final assembly per species was the assessed run with the highest *C* + *S* − 2*D* on BUSCO percentages. BUSCO v6.1 was run in genome mode (`-m genome--offline`) with 32 threads against diptera_odb12.2, embryophyta_odb12.2 and vertebrata_odb12.2 (all created 2026-05-13).

### Agreement analysis

The set of contigs retained by each RW and SD run was reconstructed from the run’s threshold triple by applying the removal rule to the alignment cache; the reconstruction reproduced the run’s output exactly for 26 of 30 runs and within one contig for the remaining 4. For each species, lineage and optimizer, the intersection and union of the sets kept under ODB9, ODB10 and ODB12.2 were computed and their Jaccard index reported, and the RW and SD intersections were compared in the same way.

### Timing benchmarks

The CPU and GPU comparison for RW and SD used *P. americana* with the embryophyta ODB9 table (5,122 contigs, 1,440 orthologs, 49,285 alignment records after prefiltering), with data loaded once and only the optimization phase timed, five replicates per configuration, 10 CPU processes against 84 GPU walkers, and total iteration budgets of 1,000, 5,000, 10,000, 50,000 and 100,000. Means, standard deviations and coefficients of variation were computed over the five replicates and speedups on the means. This benchmark was run on 2026-07-01 with an earlier ortholog table (best cost 0.0884 against 0.1036 in Table 2); the raw benchmark output was not preserved, and the timings are taken from the run summary. End-to-end wall times in Tables 2, 7 and 8 include loading of the alignment cache and ortholog table and writing of the output assemblies.

### Comparison with the original HapSolo

The original HapSolo (v0.1, Python 2.7) and a CPU-only refactor of HapSolo2 were run on *P. americana* with the embryophyta ODB9 ortholog table; the original HapSolo run used 50,000 iterations (Solares et al. 2023) and a lower bound of 0.2 on each threshold. These legacy runs were produced by earlier code and are reported as such.

### Assembly statistics

Contig statistics (contig counts at size thresholds, total length, largest sequence, N50, N75, L50, L75, NG50, NG75, LG50, LG75, GC content, N density, and the size and genome fraction of the largest *n* sequences) and cumulative length plots were produced with assembly_stats.py from the HapSolo2 package.

### Computational resources and software

All runs were made on one server with two Intel Xeon E5-2695 v2 CPUs (48 threads), 377 GB of RAM and one NVIDIA RTX A6000 GPU (48 GB, 84 streaming multiprocessors, compute capability 8.6), with Python 3.12, numpy 2.3, pandas 2.3, CuPy 14.1.1 (CUDA 12), minimap2 2.31 and miniprot 0.18. The released Apptainer image (`hapsolo2.def`) pins Python 3.12.14, pandas 2.3.3, numpy 2.3.5, CuPy 14.2.0 (`cupy-cuda12x[ctk]`, bundling the CUDA 12 runtime so that only the host driver is required), matplotlib 3.10.7, minimap2 2.31 and miniprot 0.18, and records these versions in `/opt/HapSolo/VERSIONS.txt`. BUSCO assessments of the *A. radiata* candidates were run on a separate SLURM cluster, with 32 threads per job and the same BUSCO v6.1 image and lineage dataset.

## Declarations

### Availability of data and materials

HapSolo2 is released under the Apache License 2.0 at https://github.com/esolares/HapSolo. The Apptainer definition file is in `dev/hapsolo2/hapsolo2.def`, from which the image is built with `apptainer build`. *A. funestus*: NCBI BioProject PRJNA494870 (Ghurye et al. 2019; Solares et al. 2021). *A. radiata*: Vertebrate Genomes Project assembly sAmbRad1, GenBank GCA_010909765.1 (BioProject PRJNA591369, BioSample SAMN09948523), with the raw sequence data (PacBio, Illumina and Hi-C; SRA study SRP456386) under BioProject PRJNA1008113 and at https://vgp.github.io/genomeark/Amblyraja_radiata/ (Rhie et al. 2021; Solares et al. 2021). *P. americana* cv. Gwen: NCBI BioProject PRJNA758103 (PacBio and Illumina reads and the scaffolded assembly) and Zenodo, https://doi.org/10.5281/zenodo.6392169 (Solares et al. 2023). *V. vinifera* cv. Chardonnay: NCBI BioProject PRJNA550461 (Zhou et al. 2019; Solares et al. 2021). ODB10 and ODB12.2 lineage datasets: https://busco-data.s3.amazonaws.com/lineages/. ODB9 lineage datasets: Zenodo, https://doi.org/10.5281/zenodo.20987183. The scripts that generate the tables (`analysis/build_publication_tables.py`) and the result files are in the same repository.

## Funding

This work was supported (in part) by the University of California President’s Postdoctoral Fellowship awarded to E. A. S. This work was also supported by Eurosemillas, S.A. (awarded to Mary Lu Arpaia, M. L. A.) and by the National Institute of Food and Agriculture, Agriculture and Food Research Initiative (Grant No. 2021-67013-34237, awarded to M. L. A., et al.).

## Authors’ contributions

JY prototyped GPU kernels for alignment filtering and ortholog scoring and an independent steepest-descent variant. MA and PG implemented the steepest-descent optimizer and its modular structure. EAS conceived the approach, designed the GPU implementation and the simulated annealing optimizer, wrote the package, ran the benchmarks and wrote the paper. JY, MA and PG contributed equally. All authors read and approved the manuscript.

## Acknowledgements

This work used Expanse at the San Diego Supercomputer Center (SDSC) and Bridges-2 at the Pittsburgh Supercomputing Center (PSC) through allocation MCB180035 (Species Rescue and Rapid Low Cost Assembly of Food for Structural Variant Detection using Artificial Intelligence) from the Advanced Cyberinfrastructure Coordination Ecosystem: Services & Support (ACCESS) program (Boerner et al. 2023), which is supported by U.S. National Science Foundation grants #2138259, #2138286, #2138307, #2137603 and #2138296. E. A. S. thanks Dr. Brandon Gaut for all his mentorship and help, and Dr. Pavel Pevzner for his mentorship and guidance while junior teaching faculty at UC San Diego. A special thank you to Dr. Rick Lathrop, for all his mentorship throughout my undergraduate studies until I started my position at UC San Diego in 2023, before his passing. I have finally released HapSolo2 as I promised him, and implemented simulated annealing just as he recommended several years ago. I only wish he were able to see it.

## Competing interests

The authors declare that they have no competing interests.

## Ethics approval and consent to participate

Not applicable.

## Consent for publication

Not applicable.

## Supplementary figures

**Figure S1:**
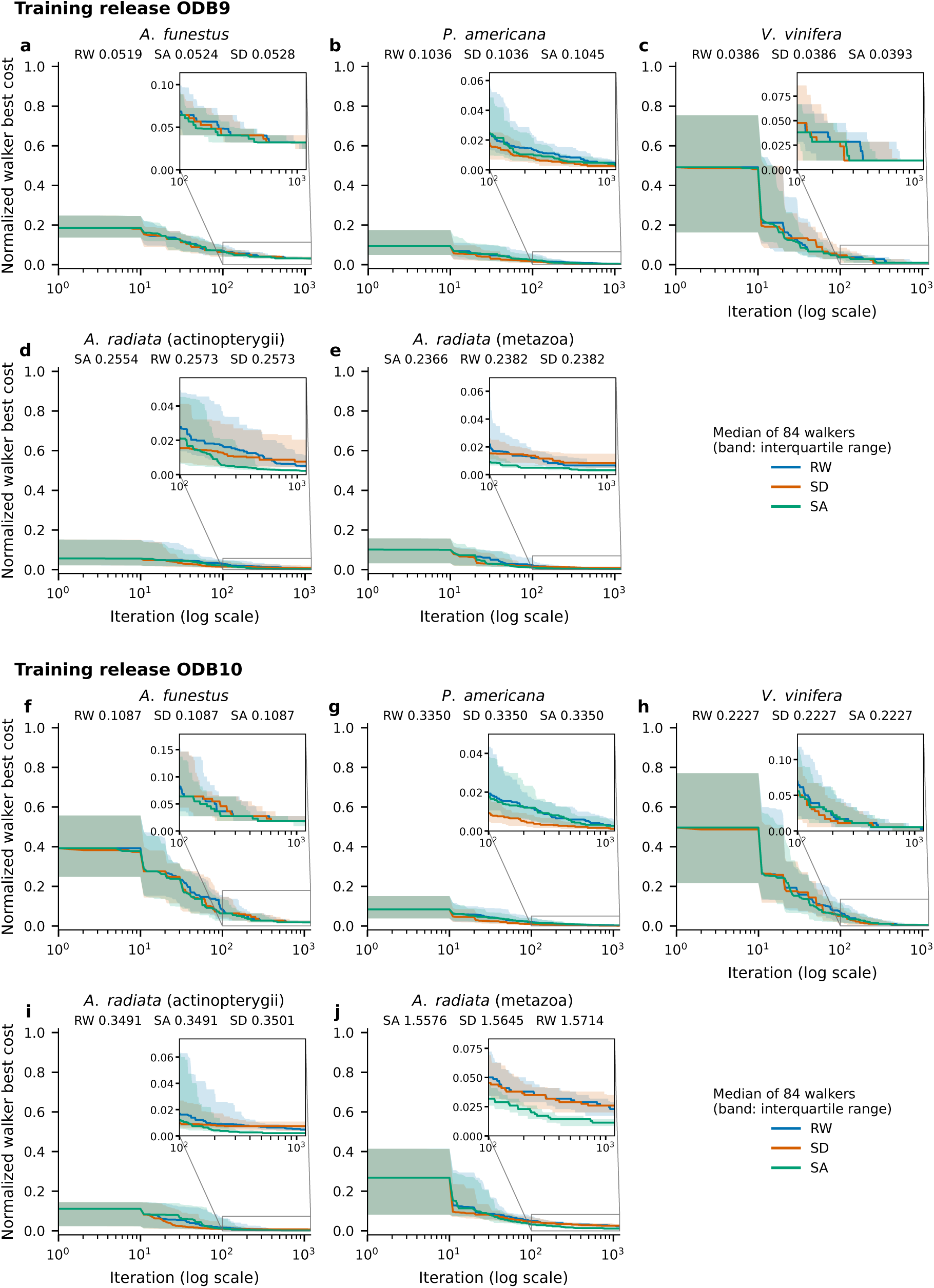
Convergence of RW, SD and SA with training releases ODB9 (a to e) and ODB10 (f to j), drawn as in Figure 2: median across 84 walkers of each walker’s best cost so far, with the interquartile range shaded, normalized per panel, and insets enlarging iterations 100 to 1,191. Panels follow the species and lineage order of Figure 2.

## Notes

https://github.com/ESB-AI-Lab/HapSolo

### Competing Interest Statement

The authors have declared no competing interest.

