## Supplementary figures and images for "HapSolo2: GPU-accelerated optimization for removing secondary haplotigs from diploid genome assemblies"

### Supplementary Figure S1

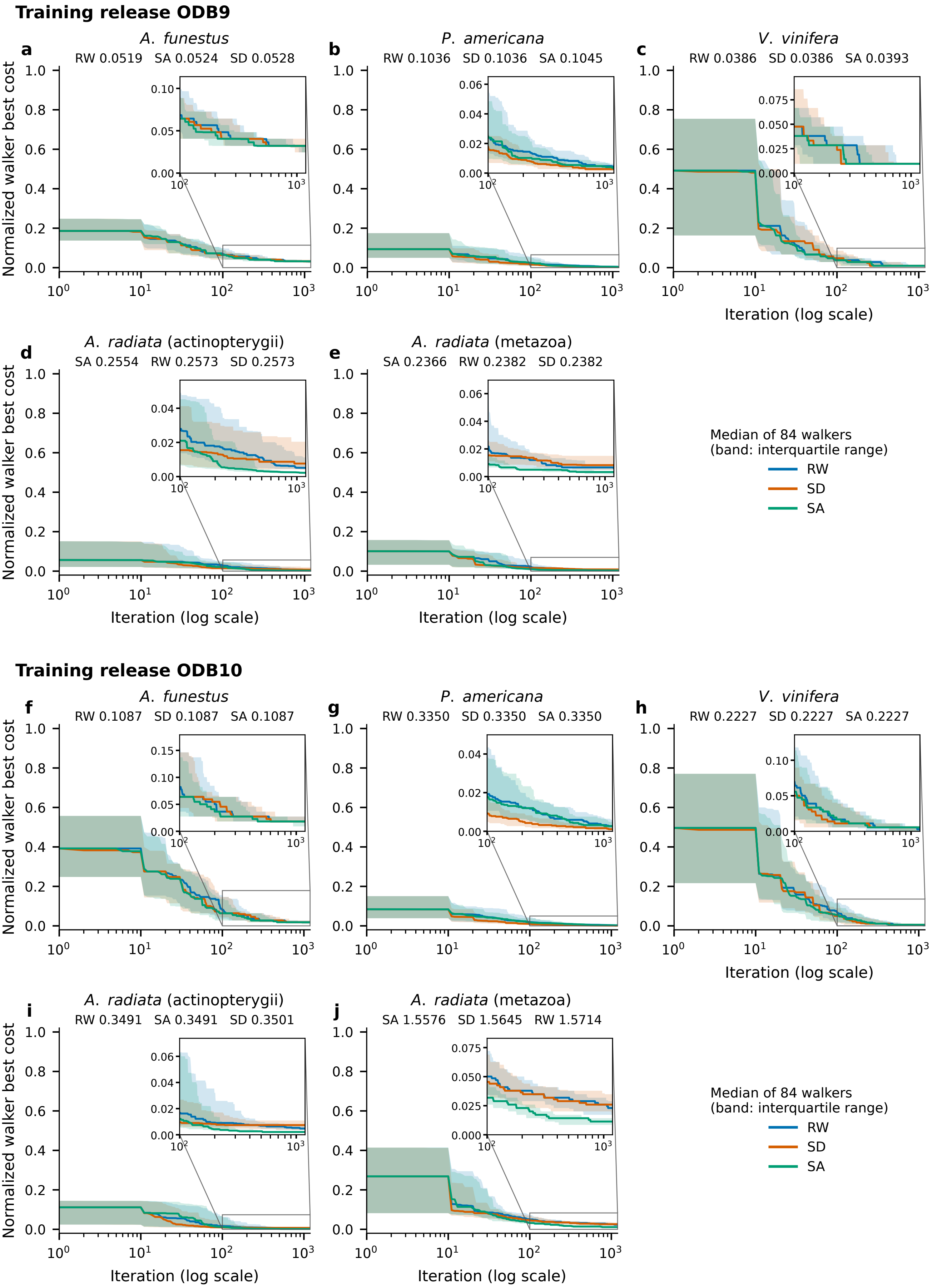
